# COPI-dependent intra-Golgi recycling at an intermediate stage of cisternal maturation

**DOI:** 10.1101/2025.09.20.677526

**Authors:** Adam H. Krahn, Areti Pantazopoulou, Jotham Austin, Natalie Johnson, Conor F. Lee-Smith, Benjamin S. Glick

## Abstract

The traffic pathways that recycle resident Golgi proteins during cisternal maturation are not completely defined. We addressed this challenge using the yeast *Saccharomyces cerevisiae*, in which maturation of individual cisternae can be visualized directly. A new assay captures a specific population of Golgi-derived vesicles at the bud neck, thereby revealing which resident Golgi proteins are carried as cargo in those vesicles. This method supplies evidence for at least three classes of intra-Golgi vesicles with different cargo compositions. Consistent with our previously published data, one class of vesicles mediates a late pathway of intra-Golgi recycling with the aid of the AP-1 and Ent5 clathrin adaptors, and a second class of vesicles mediates an early pathway of intra-Golgi recycling with the aid of the COPI vesicle coat. Here, we identify another COPI-dependent pathway of intra-Golgi recycling and show that it operates kinetically between the two previously known pathways. Thus, intra-Golgi recycling is mediated by multiple COPI-dependent pathways followed by a clathrin-dependent pathway.

## Introduction

The eukaryotic Golgi apparatus is a central hub for the modification and distribution of proteins and lipids (Farquhar and Hauri, 1997). This organelle consists of disk-like membrane-bound cisternae, which may be stacked or non-stacked depending on the organism (Mowbrey and Dacks, 2009). Two major activities occur in the Golgi. One activity is enzymatic biosynthesis and processing. Secretory proteins passing through the Golgi undergo modifications such as glycan remodeling and proteolytic cleavage, and various lipids are synthesized in the cisternal membranes. The other activity is protein sorting. Multiple classes of vesicles emerge from the Golgi to deliver material to the plasma membrane or the endolysosomal system.

These activities of the Golgi occur in a defined temporal order, and they are coordinated with biochemical changes in the cisternae (Glick and Nakano, 2009). Researchers are still seeking the best way to describe the Golgi system. A traditional approach divides the Golgi into compartments called *cis*, medial, *trans*, and *trans*-Golgi network (TGN). However, those terms have not been rigorously defined, and the compartment-focused view is arguably more of a hindrance than a help (Pantazopoulou and Glick, 2019). We favor a different approach that represents the Golgi as a set of maturing cisternae (Glick and Malhotra, 1998). In this view, each cisterna forms, undergoes a series of transformations, and ultimately disassembles, while the resident Golgi proteins continually recycle from older to younger cisternae. Membrane recycling causes a cisterna to mature—vesicles fuse with the cisterna to deliver resident Golgi proteins, and other vesicles bud from the cisterna to remove other resident Golgi proteins, so the composition of the cisterna changes over time. Presumably, a molecular logic circuit links the various membrane recycling pathways to orchestrate the sequential transformations experienced by a cisterna (Pantazopoulou and Glick, 2019). From this perspective, understanding the Golgi will require characterizing the membrane recycling pathways that drive cisternal maturation. The specific questions are: How many membrane recycling pathways operate at the Golgi? When during maturation does each pathway operate? Which traffic machinery components mediate each pathway? Which resident Golgi proteins follow each pathway?

*S. cerevisiae* is uniquely suited to answering these questions because it contains non-stacked Golgi cisternae that are optically resolvable (Wooding and Pelham, 1998). To visualize cisternal maturation, resident Golgi proteins are labeled with fluorescent tags, and changes in the composition of an individual cisterna are tracked by 4D microscopy (Losev et al., 2006; Matsuura-Tokita et al., 2006). If two or more resident Golgi proteins are labeled with different fluorophores, then their relative arrival and departure times can be measured, a method that we term “kinetic mapping”. The kinetic signature of a resident Golgi protein is useful for assigning that protein to a particular membrane recycling pathway.

A key feature of a membrane recycling pathway is the traffic machinery components that promote vesicle formation, targeting, and fusion. For intra-Golgi recycling pathways, attention has focused on vesicle formation by the COPI coat (Arab et al., 2024). COPI vesicles mediate retrograde intra-Golgi recycling of resident Golgi proteins as well as retrograde Golgi-to-ER recycling (Rabouille and Klumperman, 2005; Barlowe and Miller, 2013). It is thought that multiple COPI-dependent intra-Golgi pathways recycle different subsets of resident Golgi proteins (Sahu et al., 2022), but the number and properties of those pathways have been uncertain. Meanwhile, our studies of the yeast Golgi showed that COPI-dependent recycling is restricted to resident early Golgi proteins, and that resident late Golgi (TGN) proteins recycle with the aid of the AP-1 clathrin adaptor (Papanikou et al., 2015; Day et al., 2018; Casler et al., 2022). Similarly, a recent study of mammalian cells concluded that AP-1 mediates the recycling of late Golgi proteins (Robinson et al., 2024). In yeast, the Ent5 clathrin adaptor cooperates with AP-1 to drive intra-Golgi recycling (Casler et al., 2022). It therefore seems that recycling of resident Golgi proteins involves the actions of two vesicle budding machineries: COPI early in maturation, and clathrin together with AP-1 (plus Ent5 in yeast) late in maturation.

The question then becomes, how many distinct Golgi recycling pathways utilize each of these vesicle budding machineries? By combining kinetic mapping with functional tests, we previously suggested that two sequential AP-1/Ent5-dependent recycling pathways operate at the yeast Golgi (Casler et al., 2022). We have now revisited this issue using an updated toolkit that reveals whether different resident Golgi proteins travel together in the same vesicles. The new evidence described here points to the existence of a single AP-1/Ent5-dependent recycling pathway plus at least two COPI-dependent recycling pathways. This advance brings us closer to a complete picture of how proteins localize within the Golgi.

## Results

### Golgi-derived vesicles containing Kex2 can be captured at the yeast bud neck

We devised a method to capture Golgi-derived vesicles that contain a specific resident Golgi transmembrane protein (Fig. 1 A). The chosen capture site is the bud neck because this region is compact, easily identified, and often devoid of Golgi cisternae. Two copies of FK506-binding protein (FKBP) are fused by gene replacement to the bud-neck-localized septin Shs1 (Iwase et al., 2007), which can be tagged without visibly perturbing cytokinesis. Also present in the Shs1 fusion construct is the red fluorescent protein mScarlet (Bindels et al., 2017; Valbuena et al., 2020). The parental yeast strain has mutations that prevent growth arrest by rapamycin (Haruki et al., 2008; Papanikou et al., 2015). In addition, to facilitate the use of rapamycin and of HaloTag ligands, the parental strain lacks the transcription factors Pdr1 and Pdr3, which drive expression of pleiotropic drug transporters (Schüller et al., 2007; Barrero et al., 2016). For the capture assay, a resident Golgi transmembrane protein is tagged on its cytosolic domain with FKBP-rapamycin binding domain (FRB). Addition of rapamycin promotes heterodimerization of FKBP with FRB (Haruki et al., 2008), resulting in capture at the bud neck of vesicles containing the FRB-tagged Golgi protein. To minimize indirect effects that might alter traffic pathways, the rapamycin treatment time is only 5 min and is terminated by aldehyde fixation. If a second resident Golgi protein is tagged with GFP, then rapamycin-dependent accumulation of GFP fluorescence at the bud neck reflects co-capture of the GFP-tagged protein with the FRB-tagged protein, implying that the two proteins are in the same vesicles (Fig. 1 A).

**Figure 1.**
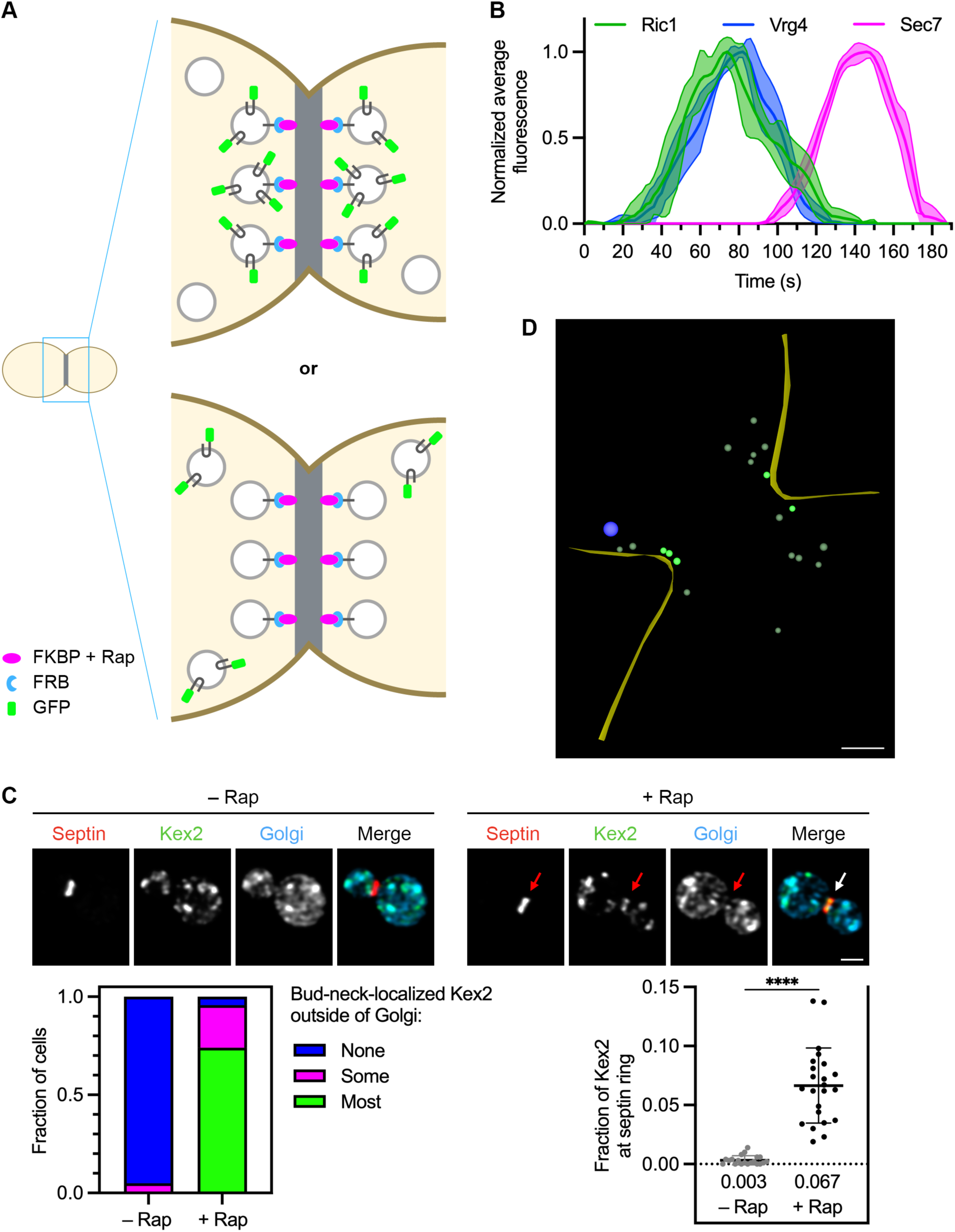
Transport vesicles carrying resident Golgi proteins can be captured at the bud neck. **(A)** Diagram of the vesicle capture assay. A septin at the bud neck is tagged with FKBP (magenta), and a resident Golgi transmembrane protein is tagged on its cytosolic domain with FRB (blue). When the resident Golgi protein is packaged into transport vesicles in the presence of rapamycin (“Rap”), some of those vesicles will be captured at the bud neck. If a second resident Golgi protein is tagged with GFP (green), then that second protein will either be co-captured at the bud neck if it travels in the same vesicles as the FRB-tagged protein (top) or not co-captured if it travels in different vesicles than the FRB-tagged protein (bottom). **(B)** Golgi maturation kinetics of GFP-tagged Ric1 compared to HaloTag-labeled Vrg4 and mScarlet-tagged Sec7. Shown are normalized and averaged traces for 11 individual cisternae. **(C)** Rapamycin-dependent capture by an FKBP-tagged septin (red) of Kex2-FRB-GFP (green). Golgi cisternae were marked by HaloTag-labeled Ric1 and Sec7 (blue). Representative images are shown. Arrows indicate non-Golgi signal at the bud neck after treatment for 5 min with rapamycin. At the lower left, capture of Kex2-FRB-GFP at the bud neck was quantified by assigning cells to categories based on whether any bud-neck-localized Kex2 signal was visible outside of Golgi cisternae, and if so, whether less than half (“Some”) or more than half (“Most”) of that signal was outside of Golgi cisternae. At the lower right, capture of Kex2-FRB-GFP at the bud neck was quantified by measuring the fraction of the total Kex2 signal within a septin mask, which had been modified by subtraction of a Golgi mask. For this type of numerical quantification, the mean capture values are displayed as thick horizontal bars and are listed numerically below the plots, and the standard deviations are displayed as thin horizontal bars. ****, significant at P value <0.0001. **(D)** Cryo-ET of vesicles captured at the bud neck using Kex2-FRB. A log-phase culture of cells expressing Kex2-FRB and Shs1-FKBP was treated with rapamycin for 5 min followed by cryopreservation and processing for cryo-ET. Shown is the model of a SIRT-reconstructed tomogram from a large budded cell. The full data set is provided in Video 1. Cell cortex, yellow; secretory vesicle, blue; putatively captured non-secretory vesicles, bright green; other non-secretory vesicles, dull green. Scale bar, 250 nm.

A limitation of this approach is that *S. cerevisiae* Golgi cisternae are mobile in the cytoplasm (Wooding and Pelham, 1998; Losev et al., 2006), so a cisterna that collides with the septin ring could be captured by the FKBP-FRB interaction, thereby generating a confounding signal due to GFP-tagged protein molecules present in the cisterna rather than in vesicles. This problem can be avoided by excluding Golgi cisternae from the analysis. For this purpose, cisternae are labeled by appending HaloTag to two Golgi proteins, Ric1 and Sec7, that should be absent from vesicles. Both proteins are peripherally membrane-associated guanine nucleotide exchange factors (Siniossoglou et al., 2000; McDonold and Fromme, 2014). Ric1 is present on early Golgi cisternae (Huh et al., 2003) while Sec7 is present on late Golgi cisternae (Rossanese et al., 1999; Losev et al., 2006), and together those two proteins are expected to mark a cisterna for most of its lifetime. This expectation was verified by three-color 4D confocal microscopy (Losev et al., 2006; Johnson and Glick, 2019). Golgi cisternae were labeled by tagging Ric1 with GFP, and by tagging the early Golgi protein Vrg4 with HaloTag conjugated to a far-red dye (Grimm et al., 2021), and by tagging Sec7 with mScarlet. Fluorescence traces for individual cisternae were smoothed, aligned, and combined to obtain averaged traces (Casler et al., 2022). The results indicated that Ric1 arrives and departs roughly in synchrony with Vrg4, while Sec7 arrives and departs later after a brief period of overlap with Ric1 (Fig. 1 B).

Quantification of vesicle capture was performed as follows. A mask was created from the red fluorescence of tagged Shs1, and this septin mask was modified by subtracting a Golgi mask created from the far-red fluorescence of HaloTag-labeled Ric1 and Sec7, thereby ensuring that the septin mask excluded any cisternae at the bud neck. The Golgi mask was defined generously to encompass nearly all of the visible HaloTag signal, based on the rationale that eliminating the contribution from Golgi cisternae was more important than preserving the full signal from the captured vesicles. The criterion for analyzing a cell was that subtraction of the Golgi mask removed less than 65% of the septin mask. For the measurement, the green signal from the GFP-tagged Golgi protein was examined. The green signal within the subtraction-modified septin mask was compared with the total cellular green signal to determine the fraction of the GFP-tagged Golgi protein present in captured vesicles.

As a proof of principle, the late Golgi protein Kex2 (Fuller et al., 1988) was tagged on its cytosolic C-terminus with both FRB and GFP so that capture of Kex2-containing vesicles could be visualized. Kex2 and the other Golgi proteins used in this study were tagged by chromosomal replacement of the endogenous genes with fusion genes under control of the normal promoters. In the absence of rapamycin, tagged Kex2 was found in punctate Golgi cisternae and was usually undetectable at the bud neck (Fig. 1 C, upper left). By contrast, after incubation with rapamycin, Kex2 fluorescence was usually visible not only in punctate Golgi cisternae, but also at the bud neck outside of Golgi cisternae (Fig. 1 C, upper right). This capture of Kex2 was quantified by two methods. For the numerical quantification method, the Kex2 fluorescence that overlapped with the subtraction-modified septin mask was measured as described above. A rapamycin-dependent signal was consistently observed (Fig. 1 C, lower right). Quantification yielded average values of 6.7% of the total Kex2 fluorescence at the bud neck for rapamycin-treated cells versus 0.3% for untreated cells. Such measurements may underestimate the efficiency of rapamycin-dependent capture due to the generous Golgi masks used for subtraction, but we are confident that only a small fraction of the resident Golgi protein molecules were captured at the bud neck, implying that Golgi function was likely unperturbed. For the categorical quantification method, cells were assigned to categories based on visual estimates of how much of the Kex2 fluorescence at the bud neck was outside of Golgi cisternae. Of the rapamycin-treated cells, 96% showed bud-neck-localized Kex2 fluorescence outside of Golgi cisternae, including 74% of the cells for which a majority of the bud-neck-localized fluorescence was outside of cisternae and 22% of the cells for which a minority of the bud-neck-localized fluorescence was outside of cisternae (Fig. 1 C, lower left). Of the untreated cells, only 5% showed bud-neck-localized Kex2 fluorescence outside of cisternae. Thus, the categorical quantification is useful for verifying that results from the numerical quantification reflect effects that are evident by visual inspection. These two methods provide complementary ways to assess the capture of a fluorescent Golgi protein at the bud neck.

To confirm that the Kex2 fluorescence at the bud neck represented vesicles, we performed cryo-electron tomography (cryo-ET; Gan et al., 2019) (Fig. 1 D). Cells expressing Kex2-FRB were examined with or without rapamycin treatment under the conditions of the vesicle capture assay. We analyzed 6 untreated large budded cells and 10 rapamycin-treated large budded cells. Lamellae of thickness ∼200-220 nm, representing an estimated 15-30% of the total bud neck volume (Bertin et al., 2012), were imaged and modeled. With or without rapamycin treatment, electron-dense vesicles of diameter ∼90 nm were occasionally seen near the bud neck. Those structures were presumably secretory vesicles because they resembled the secretory vesicles clustered in the buds of small budded cells (data not shown). Secretory vesicles were expected to be present near the bud neck of large budded cells (Finger et al., 1998; Pruyne et al., 2004). Also visible were less electron-dense non-secretory vesicles of diameters ∼35-60 nm. To score non-secretory vesicles that had putatively been captured by the FKBP-tagged septin, we used prior ultrastructural evidence that yeast septin rings extend about 200 nm in each direction along the cell cortex from the center of the bud neck (Bertin et al., 2012; Ong et al., 2014). Based on that number and the predicted lengths of the FKBP-tagged Shs1 septin (48 nm) and the FRB-tagged Kex2 cytosolic tail (58 nm), we defined capture zones for the bud neck. The boundaries of a capture zone were determined by (a) extending 200 nm in each direction along the cell cortex from the center of the bud neck, and then (b) from the resulting region of plasma membrane, extending 106 nm into the cytoplasm. A vesicle was counted as putatively captured if any part of its membrane was within a capture zone. For the non-rapamycin-treated cells, the number of vesicles meeting this criterion ranged from 0 to 2 (mean = 0.7). For the rapamycin-treated cells, the number of putatively captured vesicles ranged from 0 to 7 (mean = 2.6). Such variability was anticipated based on the variable capture signals seen by fluorescence imaging. Fig. 1 D and Video 1 show an example of a rapamycin-treated cell for which the lamella included 5 putatively captured vesicles (marked in bright green). Thus, the combined fluorescence and cryo-ET data indicate that our assay is suitable for capturing vesicles that contain a specific resident Golgi transmembrane protein.

### Two Golgi proteins that have the same recycling kinetics as Kex2 are present in Kex2-containing vesicles

We previously showed that Kex2 recycles within the Golgi in a pathway dependent on the AP-1 and Ent5 clathrin adaptors (Casler et al., 2022). Other late Golgi proteins, including Ste13 and Stv1 (Table 1), were proposed to follow the same recycling pathway because they closely resembled Kex2 in their kinetics of arrival and departure during Golgi maturation (Casler et al., 2022). By contrast, the early Golgi protein Vrg4 (Table 1) recycles in a COPI-dependent manner (Abe et al., 2004; Mari et al., 2014), and it arrives and departs much sooner than Kex2 (Papanikou et al., 2015). The prediction was therefore that Ste13 and Stv1 should be co-captured in Kex2-containing vesicles whereas Vrg4 should not.

**Table 1.** Resident Golgi transmembrane proteins examined in this study. The proteins marked in bold are proposed as reference markers for the three intra-Golgi recycling pathways described here. The proteins marked with an asterisk (“*”) were tagged with FRB for use in the vesicle capture assay.

| Resident Golgi transmembrane protein | Inferred major recycling pathway(s) |
| --- | --- |
| **Kex2\*** | **AP-1/Ent5** |
| Ste13 | AP-1/Ent5 |
| Stv1 | AP-1/Ent5 |
| Sys1* | COPI(b') + AP-1/Ent5 ‡ |
| Aur1 | COPI(b') + AP-1/Ent5 ‡ |
| **Tmn1** | **COPI(b')** |
| Gnt1 | COPI(b') |
| **Vrg4\*** | **COPI(b)** |
| Kre2 | COPI(b) |
| Pmr1 | COPI(b) |

‡ It is possible that native Sys1 and Aur1 traffic mainly in the COPI(b') pathway and that the fluorescent tags divert a fraction of the molecules into the AP-1/Ent5 pathway.

To test this prediction, Kex2 was tagged with FRB alone, and either Ste13, Stv1, or Vrg4 was tagged with GFP. After rapamycin addition, Ste13 was seen to accumulate at the bud neck (Fig. 2 A). The pattern of accumulation was variable—in some cells the Ste13 signal extended across the bud neck, and in other cells the Ste13 signal was concentrated in one part of the bud neck (Fig. S1 A, top two rows). We speculate that passage of the nucleus into the daughter cell can affect which parts of the bud neck are accessible for vesicle capture. As judged by the categorical quantification method, 88% of the rapamycin-treated cells showed bud-neck-localized Ste13 with most or some of the signal outside of Golgi cisternae (Fig. 2 A). Examples of cells that were assigned to different categories are shown in Fig. S1 A. As judged by the numerical quantification method, an average of 3.6% of the cellular Ste13 fluorescence was at the bud neck in rapamycin-treated cells versus 0.3% in untreated cells (Fig. 2 A). For Stv1, similar results were obtained (Fig. 2 B and Fig. S1 B). For Vrg4, as expected, the fluorescence at the bud neck outside of Golgi cisternae was very low both with and without rapamycin addition (Fig. 2 C and Fig. S1 C). Thus, capture of vesicles containing FRB-tagged Kex2 leads to selective co-capture of Ste13 and Stv1.

**Figure 2.**
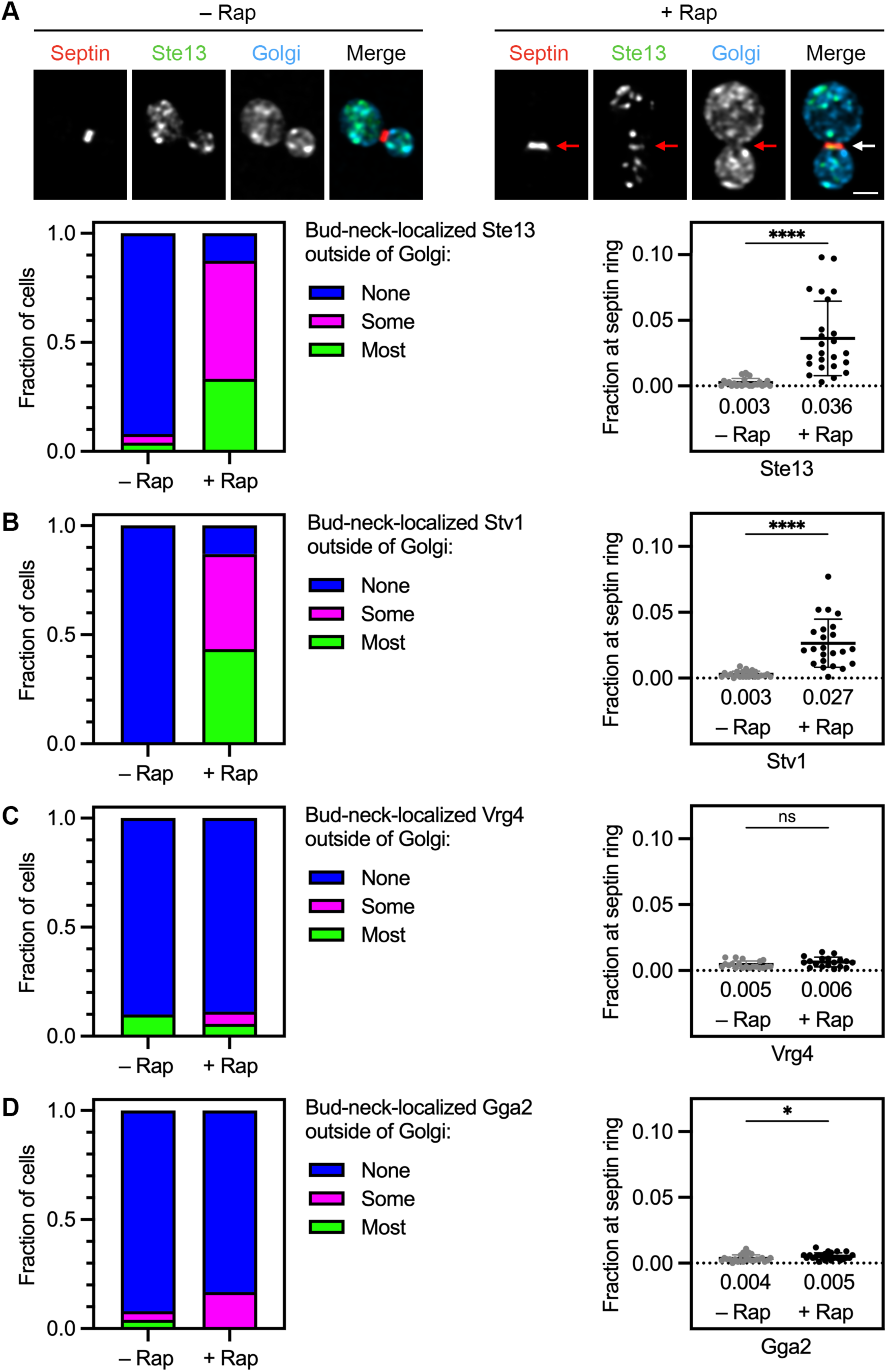
AP-1/Ent5 cargoes can be specifically captured at the bud neck together with Kex2. **(A)** Rapamycin-dependent capture with Kex2-FRB of GFP-tagged Ste13 (green) by an FKBP-tagged septin (red). Golgi cisternae were marked by HaloTag-labeled Ric1 and Sec7 (blue). Representative images are shown. Arrows indicate non-Golgi signal at the bud neck in cells treated for 5 min with rapamycin (“Rap”). At the lower left, capture of Ste13 at the bud neck was quantified by assigning cells to categories as described in Fig. 1 C. At the lower right, capture of Ste13 at the bud neck was quantified numerically as described in Fig. 1 C. ****, significant at P value <0.0001. **(B)** Capture with Kex2-FRB of GFP-tagged Stv1. Categorical and numerical quantification was performed as in (A). ****, significant at P value <0.0001. **(C)** No capture with Kex2-FRB of GFP-tagged Vrg4. Categorical and numerical quantification was performed as in (A). ns, not significant. **(D)** Minimal capture with Kex2-FRB of GFP-tagged Gga2. Categorical and numerical quantification was performed as in (A). *, significant at P value 0.04.

As described above, subtraction of the Golgi mask ensured that with FRB-tagged Kex2, the Ste13 and Stv1 signals resulted from capture of vesicles rather than from capture of late Golgi cisternae at the bud neck. To confirm the effectiveness of this method, we marked late Golgi cisternae with the clathrin adaptor Gga2, which has a kinetic signature largely overlapping that of Kex2 (Daboussi et al., 2012; Casler and Glick, 2020). Gga2 mediates transport from the Golgi to prevacuolar endosomes and should be absent from Kex2-containing AP-1/Ent5 vesicles (Myers and Payne, 2013). For Gga2, the fluorescence at the bud neck outside of Golgi cisternae was very low as judged by both of the quantification methods (Fig. 2 D and Fig. S1 D), indicating that the subtraction-modified septin masks enable measurement of fluorescence from captured vesicles while excluding nearly all of the fluorescence from Golgi cisternae. This control experiment reinforces the conclusion that Ste13 and Stv1 travel together with Kex2 in vesicles that are largely devoid of Vrg4.

### Sys1 and Aur1 arrive at maturing cisternae before Kex2 but are nevertheless found in Kex2-containing vesicles

We previously reported that the resident Golgi transmembrane proteins Sys1 and Aur1 had kinetic signatures between those of Vrg4, which is COPI-dependent, and Kex2, which is AP-1/Ent5-dependent (Casler et al., 2022). The assumption was that Sys1 and Aur1 defined a distinct intra-Golgi recycling pathway. To test this idea, we began by extending the earlier kinetic analysis of Sys1. Cells expressing Halo-tagged Sys1 and GFP-tagged Kex2 were analyzed by 4D confocal microscopy. A representative example is shown in Fig. S2 A and Video 2. Sys1 typically began to arrive at a cisterna about 20 s before Kex2. Then Sys1 typically began to depart from the cisterna sooner than Kex2, although some Sys1 signal remained detectable until Kex2 had almost completely departed. Averaged two-color fluorescence traces are shown in Fig. 3 A (left). In a separate experiment, GFP-tagged Aur1 was tracked together with Halo-tagged Sys1. The results confirmed that the kinetic signature of Aur1 is very similar to that of Sys1 (Fig. 3 A, right). Because Sys1 and Aur1 begin to arrive before Kex2, we infer that they travel at least some of the time in vesicles lacking Kex2.

**Figure 3.**
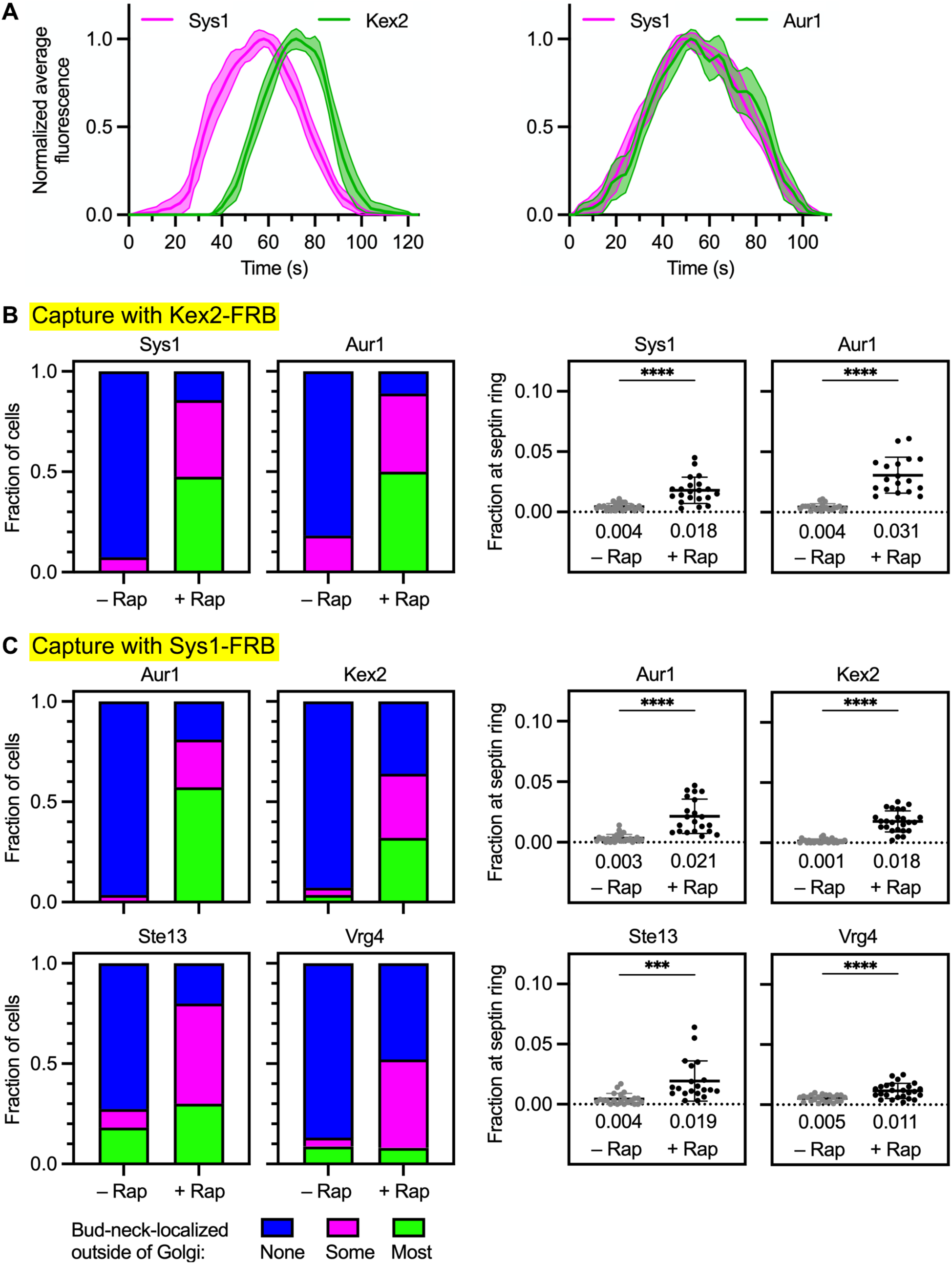
Sys1 and Aur1 are present in AP-1/Ent5 vesicles. **(A)** Golgi maturation kinetics of HaloTag-labeled Sys1 compared to GFP-tagged Kex2 or Aur1. Shown are normalized and averaged traces for 11 individual cisternae for the Sys1/Kex2 comparison and 10 individual cisternae for the Sys1/Aur1 comparison. **(B)** Capture with Kex2-FRB of GFP-tagged Sys1 and Aur1. Categorical and numerical quantifications were performed as in Fig. 2 A. ****, significant at P value <0.0001. **(C)** Capture with Sys1-FRB of GFP-tagged Aur1 and Kex2 and Ste13, and weak capture with Sys1-FRB of GFP-tagged Vrg4. Categorical and numerical quantifications were performed as in Fig. 2 A. ****, significant at P value <0.0001; ***, significant at P value 0.0008.

The assumption that Sys1 and Aur1 follow a distinct recycling pathway led us to predict that neither Sys1 nor Aur1 should be co-captured in vesicles containing FRB-tagged Kex2. But surprisingly, both proteins were co-captured with FRB-tagged Kex2 (Fig. 3 B and Fig. S2 B). A possible explanation is that Sys1 and Aur1 partition between two pathways: a pathway with intermediate kinetics of both arrival and departure, and the AP-1/Ent5 pathway followed by Kex2. This hypothesis could explain the relative kinetic signatures of Sys1 and Kex2, according to the following scenario. Sys1 arrival occurs in two phases, the first one earlier than Kex2 arrival and the second one synchronous with Kex2 arrival, and then Sys1 departure occurs in two phases, the first one earlier than Kex2 departure and the second one synchronous with Kex2 departure. In this view, the empirical kinetic signature of Sys1 or Aur1 represents the sum of the kinetic signatures for two different recycling pathways.

As an initial test of the new hypothesis, we established a capture assay in which Sys1 was tagged on its cytosolic C-terminus with FRB. To verify that Sys1-containing vesicles could be captured, Sys1 was tagged with both FRB and GFP. Upon rapamycin addition, the dual-tagged Sys1 accumulated in vesicles at the bud neck as judged by fluorescence imaging (Fig. S2 C). Cryo-ET confirmed that capture of Sys1-FRB resulted in the appearance of vesicles at the bud neck (Fig. S2 D and Video 3). In cells expressing Sys1 tagged with FRB alone, either Aur1, Kex2, Ste13, or Vrg4 was tagged with GFP. After rapamycin addition, Aur1 showed robust accumulation at the bud neck (Fig. 3 C and Fig. S2 E), confirming that Aur1 travels in vesicles together with Sys1. For Kex2 and Ste13, co-capture with Sys1 was also readily detectable (Fig. 3 C and Fig. S2 E). When this result is viewed in light of the kinetic data, it supports the idea that a fraction of the Sys1 molecules recycle in an intermediate pathway while the rest of the Sys1 molecules recycle in a late AP-1/Ent5-dependent pathway together with Kex2 and Ste13.

Interestingly, Vrg4 showed a weak but statistically significant signal for co-capture with Sys1 (Fig. 3 C). This effect was seen in both of the quantitative assays, even though it was hard to discern by visual examination of the micrographs (Fig. S2 E). We speculate that the weak Vrg4 signal reflects imperfect fidelity of Golgi traffic pathways. Sys1 is present in a Golgi cisterna when Vrg4 is departing (Fig. S3 A), so Sys1 molecules might occasionally be packaged into the same early pathway vesicles that recycle Vrg4. Alternatively, Vrg4 molecules might occasionally fail to recycle in their “normal” early pathway, in which case they could be salvaged by recycling together with Sys1 in the intermediate pathway. Either event could generate a signal for Vrg4 in the assay for co-capture with Sys1. The ability of the vesicle capture method to detect these types of low-frequency events must be kept in mind when seeking to identify the major recycling pathway(s) followed by a given Golgi protein. To interpret the co-capture data, we rely on comparative quantification to distinguish between relatively strong versus weak signals, and we examine complementary data from kinetic analysis.

### Tmn1 and Gnt1 follow an intermediate recycling pathway and are largely absent from Kex2-containing vesicles

Based on the evidence that Sys1 and Aur1 partition between the intermediate and late recycling pathways, we looked for proteins that recycle almost exclusively in the intermediate pathway. The evaluation was based on kinetic analysis. A candidate was Tmn1, also known as Emp70 (Singer-Krüger et al., 1993; Schimmöller et al., 1998; Woo et al., 2015). This protein is a member of the evolutionarily conserved but functionally uncharacterized “transmembrane nine” family of Golgi proteins. Tmn1 contains a C-terminal cytosolic tail that reportedly confers COPI-dependent Golgi localization (Woo et al., 2015), so the luminal N-terminus was tagged with GFP. The Ost1 signal sequence was used to ensure efficient co-translational translocation of the GFP tag (Fitzgerald and Glick, 2014). Tmn1 began to arrive after Vrg4 and finished departing after Vrg4 (Fig. 4 A, upper left), and Tmn1 began to arrive before Kex2 and finished departing before Kex2 (Fig. 4 A, upper right). Thus, Tmn1 resembled Sys1 in arriving after Vrg4 and before Kex2 (Fig. S3 A and Fig. 3 A, left). When Tmn1 and Sys1 were directly compared, the relative arrival and departure times showed significant variability (Fig. S3 B), but Sys1 typically became detectable shortly after Tmn1 and persisted significantly longer than Tmn1 (Fig. 4 A, lower left). These results fit with the idea that Tmn1 recycles in an intermediate pathway together with a fraction of the Sys1 molecules, and that Tmn1 is largely absent from the late pathway followed by Kex2 and the rest of the Sys1 molecules.

**Figure 4.**
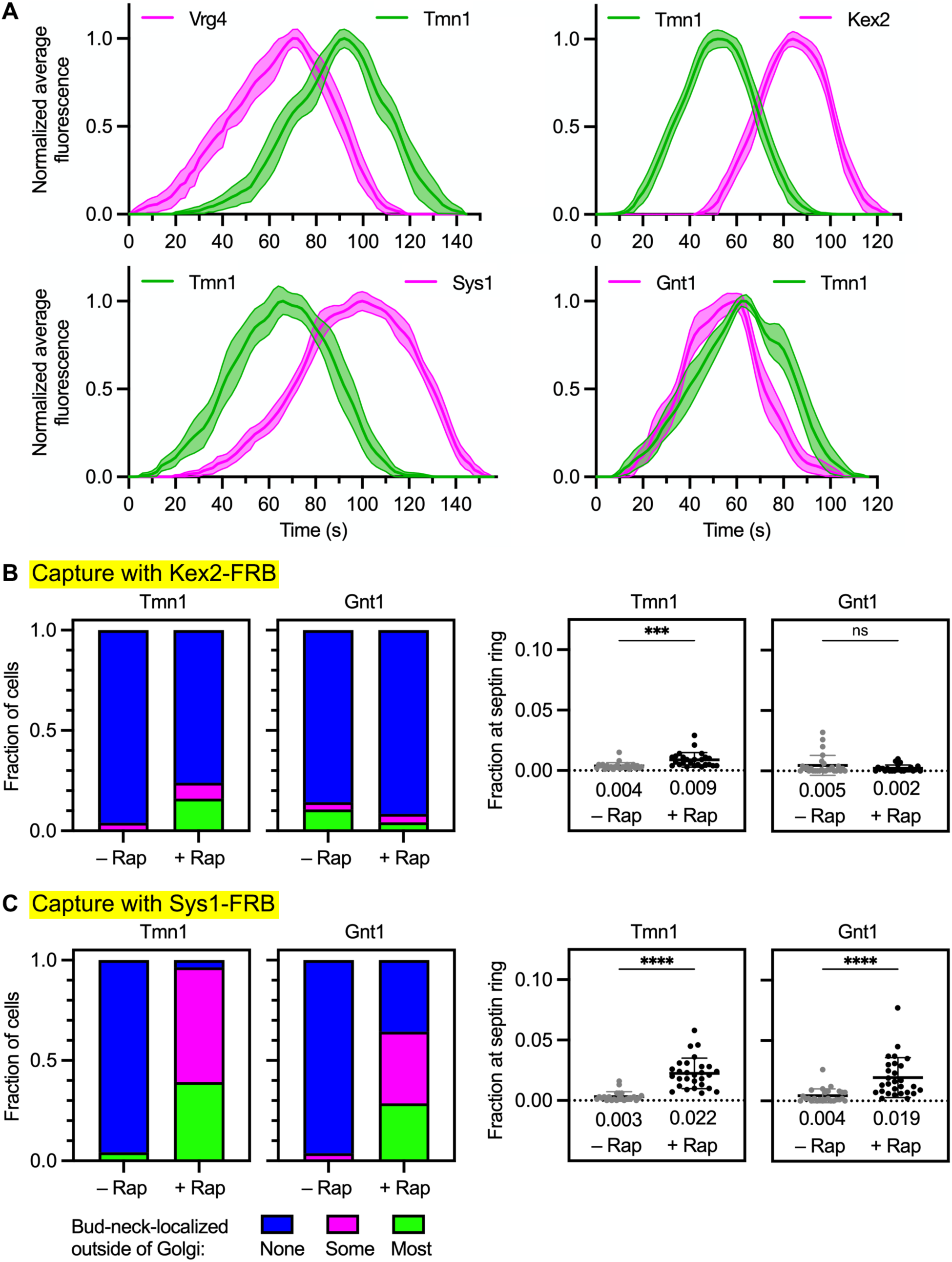
Tmn1 and Gnt1 follow an intermediate recycling pathway that is also followed by a fraction of the Sys1 molecules. **(A)** Golgi maturation kinetics of GFP-tagged Tmn1 compared to HaloTag-labeled Vrg4, Kex2, Sys1, or Gnt1. Shown are normalized and averaged traces for the following numbers of individual cisternae: 13 for the Tmn1/Vrg4 comparison, 15 for the Tmn1/Kex2 comparison, 11 for the Tmn1/Sys1 comparison, and 14 for the Tmn1/Gnt1 comparison. The strains carried the *vps10-104* mutation due to the luminal tags on Tmn1 and Gnt1. **(B)** Weak capture with Kex2-FRB of Tmn1, and no capture with Kex2-FRB of Gnt1. Categorical and numerical quantifications were performed as in Fig. 2 A. ***, significant at P value 0.0005; ns, not significant. **(C)** Capture with Sys1-FRB of Tmn1 and Gnt1. Categorical and numerical quantifications were performed as in Fig. 2 A. ****, significant at P value < 0.0001.

In addition to Tmn1, we identified Gnt1 as a candidate cargo of the intermediate pathway. Gnt1 is a type II transmembrane protein that functions as an *N*-acetylglucosaminyltransferase, and it was reported to recycle with intermediate kinetics (Yoko-o et al., 2003; Tojima et al., 2019). Gnt1 was tagged on its luminal C-terminus with HaloTag. When Gnt1 and Tmn1 were directly compared, the two proteins typically began to arrive at about the same time, but for most of the cisternae examined, Gnt1 finished departing earlier than Tmn1 (Fig. 4 A, lower right and Fig. S3 C). Thus, our kinetic analysis of Sys1, Tmn1, and Gnt1 indicates that for different cargoes of the putative intermediate recycling pathway, the arrival and departure times are somewhat variable. As explored in the Discussion, such variability can potentially be reconciled with the concept of a single molecularly defined pathway. We favor the interpretation that Tmn1 and Gnt1 recycle with each other and with a fraction of the Sys1 molecules.

To test whether Tmn1 and Gnt1 primarily follow an intermediate recycling pathway, we would ideally tag one of those proteins with FRB for use in a new vesicle capture assay. Unfortunately, neither Tmn1 nor Gnt1 can be cytosolically tagged without disrupting Golgi localization. As an alternative, we used the existing vesicle capture assays to ask whether Tmn1 and Gnt1 are present in Sys1-containing vesicles but not in Kex2-containing vesicles. In assays for co-capture with Kex2, Tmn1 showed a very weak signal and Gnt1 showed no signal (Fig. 4 B and Fig. S3 D). By contrast, in assays for co-capture with Sys1, both Tmn1 and Gnt1 showed relatively strong signals (Fig. 4 C and Fig. S3 E). The combined data support the idea that Tmn1 and Gnt1 follow an intermediate recycling pathway, that Kex2 follows a late recycling pathway, and that Sys1 partitions between those two pathways.

### The intermediate recycling pathway does not depend directly on the AP-1 and Ent5 adaptors

We previously concluded that recycling of Sys1 and Aur1 involved AP-1 and Ent5 (Casler et al., 2022). That conclusion was based on examination of a strain lacking the AP-1 subunit Apl4 as well as Ent5. In *apl4Δ ent5Δ* cells, proteins that would normally recycle within the Golgi in an AP-1/Ent5-dependent manner are expected to travel to the plasma membrane in secretory vesicles and then return to the late Golgi in endocytic vesicles (Day et al., 2018). When endocytosis is inhibited in *apl4Δ ent5Δ* cells using the Arp2/3 inhibitor CK-666, exocytosis is also inhibited, and proteins cycling between the Golgi and the plasma membrane become trapped at secretion sites either in secretory vesicles or in the plasma membrane (Casler et al., 2022). During our original analysis of the effects of CK-666 in *apl4Δ ent5Δ* cells, Kex2 and Ste13 showed strong effects, Vrg4 showed no effect, and Sys1 and Aur1 showed moderate effects (Casler et al., 2022). In hindsight, the moderate effects can be explained if the AP-1/Ent5-dependent late recycling pathway carries only a fraction of the Sys1 and Aur1 molecules.

For the intermediate recycling pathway that carries the remaining fraction of the Sys1 and Aur1 molecules together with Tmn1 and Gnt1, we predicted no dependency on AP-1/Ent5. This prediction came from observing that Tmn1 finished departing from a maturing cisterna around the time that the AP-1 subunit Apl2 began to arrive (Fig. 5 A). Because Ent5 begins to arrive simultaneously with Apl2 (Daboussi et al., 2012; Casler et al., 2022), neither AP-1 nor Ent5 is present at the Golgi during the departure phase of Tmn1, implying that Tmn1 recycling is independent of AP-1/Ent5. To verify this conclusion, we updated the assay in which *apl4Δ ent5Δ* mutant cells are treated with CK-666 (Casler et al., 2022). Cells were assigned to three categories based on whether a GFP-tagged Golgi protein was concentrated primarily in Golgi-like puncta distributed throughout the cell with no concentration at secretion sites, or whether it was concentrated primarily at secretion sites, or whether it showed a hybrid pattern of concentration both in Golgi-like puncta and at secretion sites. For example, Kex2 mainly showed concentration at Golgi-like puncta in untreated or CK-666-treated wild-type cells, and mainly showed concentration at secretion sites in CK-666-treated mutant cells, and mainly showed a hybrid pattern in untreated mutant cells (Fig. 5 B). The hybrid pattern in untreated mutant cells presumably indicates that Kex2 was cycling between the Golgi and the plasma membrane rather than recycling within the Golgi (Casler et al., 2022). Like Kex2, Ste13 showed concentration at secretion sites in CK-666-treated mutant cells (Fig. 5 B). Sys1 also showed concentration at secretion sites in CK-666-treated mutant cells, but as previously seen (Casler et al., 2022), this effect was weaker than for Kex2 and Ste13 (Fig. 5 B). A likely explanation is that only a fraction of the Sys1 molecules were missorted into secretory vesicles in *apl4Δ ent5Δ* mutant cells. Vrg4 showed no concentration at secretion sites in CK-666-treated mutant cells, as expected (Fig. 5 B). Tmn1 showed some accumulation in the vacuole in *apl4Δ ent5Δ* mutant cells with or without CK-666, perhaps due to perturbed Golgi function, but it showed minimal concentration at secretion sites in CK-666-treated mutant cells (Fig. 5 B). We conclude that the primary recycling pathway of Tmn1 is independent of AP-1/Ent5.

**Figure 5.**
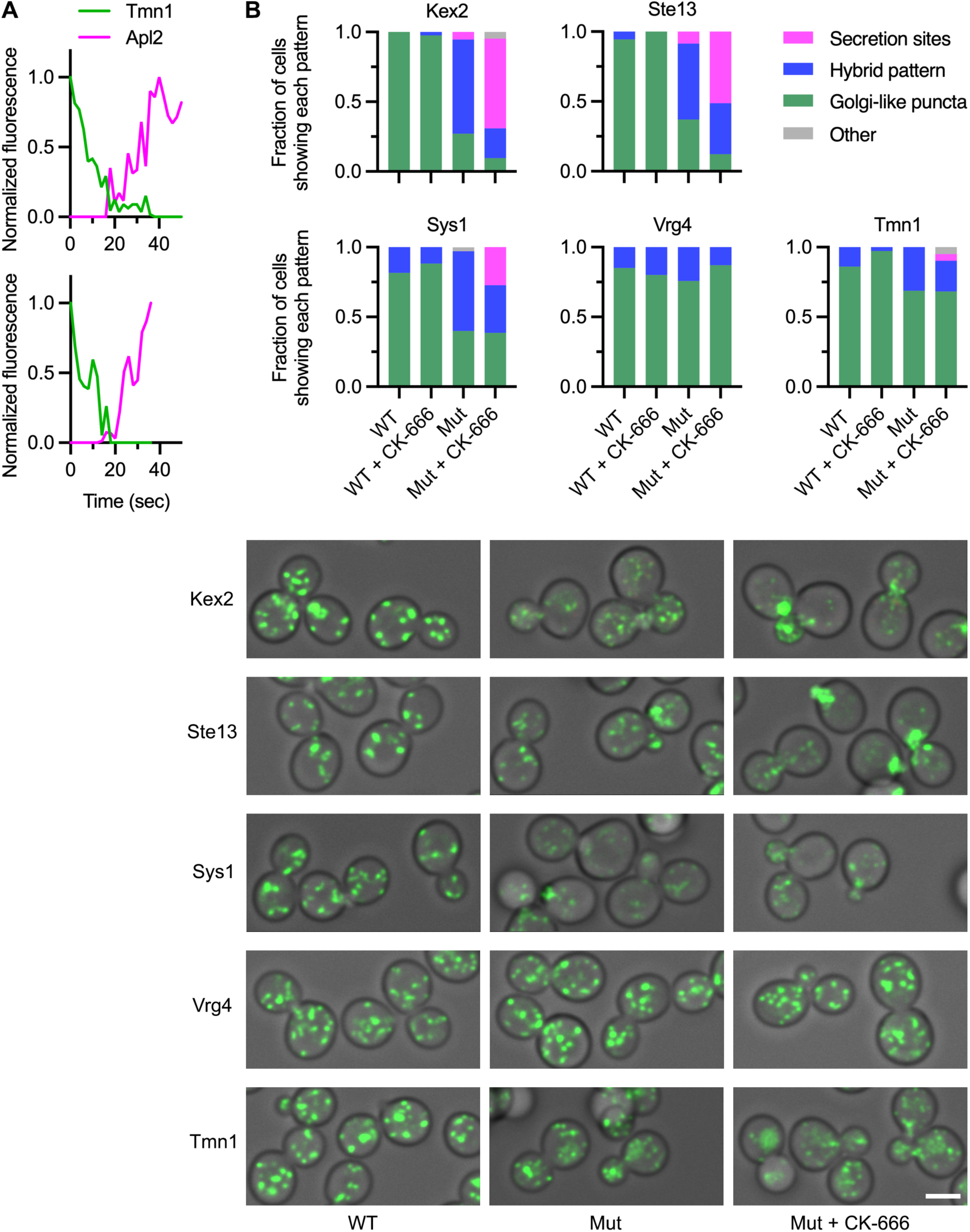
The intermediate recycling pathway does not depend on AP-1/Ent5. **(A)** Partial kinetic traces for two representative cisternae illustrating that the AP-1 subunit Apl2 arrives after Tmn1 has largely departed. For two cisternae that showed detectable kinetic overlap between GFP-tagged Tmn1 and HaloTag-labeled Apl2, the fluorescence signals were quantified for 10-15 s before and after the overlap period. The maximal value obtained for each signal was set to 1.0. **(B)** Plots and representative images showing that in cells lacking AP-1 and Ent5, inhibition of endocytosis with CK-666 affected the localizations of some Golgi proteins but not of Vrg4 or Tmn1. Wild-type “WT” or *apl4Δ ent5Δ* mutant (“Mut”) cells expressing the indicated GFP-tagged Golgi proteins were examined by fluorescence microscopy after either mock treatment or treatment for 15-25 min with CK-666. The fluorescence pattern of each tagged protein was quantified by assigning cells to categories based on whether most of the protein was in Golgi-like puncta with no visible signal at secretion sites, or whether most of the protein was concentrated at secretion sites, or whether the protein was distributed in a hybrid pattern between Golgi-like puncta and secretion sites. Approximately 30-45 cells were examined for each Golgi protein and each condition. In a few cases (“Other”), a cell could not be confidently assigned to any of the three categories. Representative images are shown for mock-treated wild-type cells, mock-treated mutant cells, and CK-666-treated mutant cells. For a given GFP-tagged protein, the images were all captured and processed with the same parameters, except that during deconvolution of the Sys1 images, a higher background subtraction was performed for WT than mutant cells.

### The intermediate recycling pathway depends on COPI

It seemed possible that the intermediate recycling pathway employs COPI. If so, then COPI should be present on Golgi cisternae throughout the departure phase of Tmn1. To test this prediction, we first used three-color kinetic mapping to compare the COPI subunit Sec26 to the early and late Golgi reference proteins Vrg4 and Sec7 (Losev et al., 2006; Papanikou et al., 2015). Previous studies indicated that COPI associates with early Golgi cisternae and persists into the arrival phase of Sec7 (Papanikou et al., 2015; Kim et al., 2016; Tojima et al., 2019). Indeed, Sec26 began to arrive on maturing cisternae well before Vrg4, and Sec26 finished departing around the time that Sec7 reached its peak abundance (Fig. 6 A). Kinetic mapping of Sec26 was technically difficult because the earliest COPI-containing cisternae were small and numerous, but the averaged data suggest that COPI may arrive at cisternae in two successive waves. For our purposes, the key point was that Sec26 persisted after Vrg4 had departed, suggesting that COPI was still present during the departure of proteins that follow the intermediate recycling pathway. Indeed, tracking of Sec26 during the residence period of Tmn1 indicated that COPI persisted at a maturing cisterna until Tmn1 had finished departing (Fig. 6 B). Of the 13 individual cisternae examined in this way, all of them showed Sec26 persisting at least as long as Tmn1. Thus, COPI could be involved in forming the vesicles of the intermediate recycling pathway.

**Figure 6.**
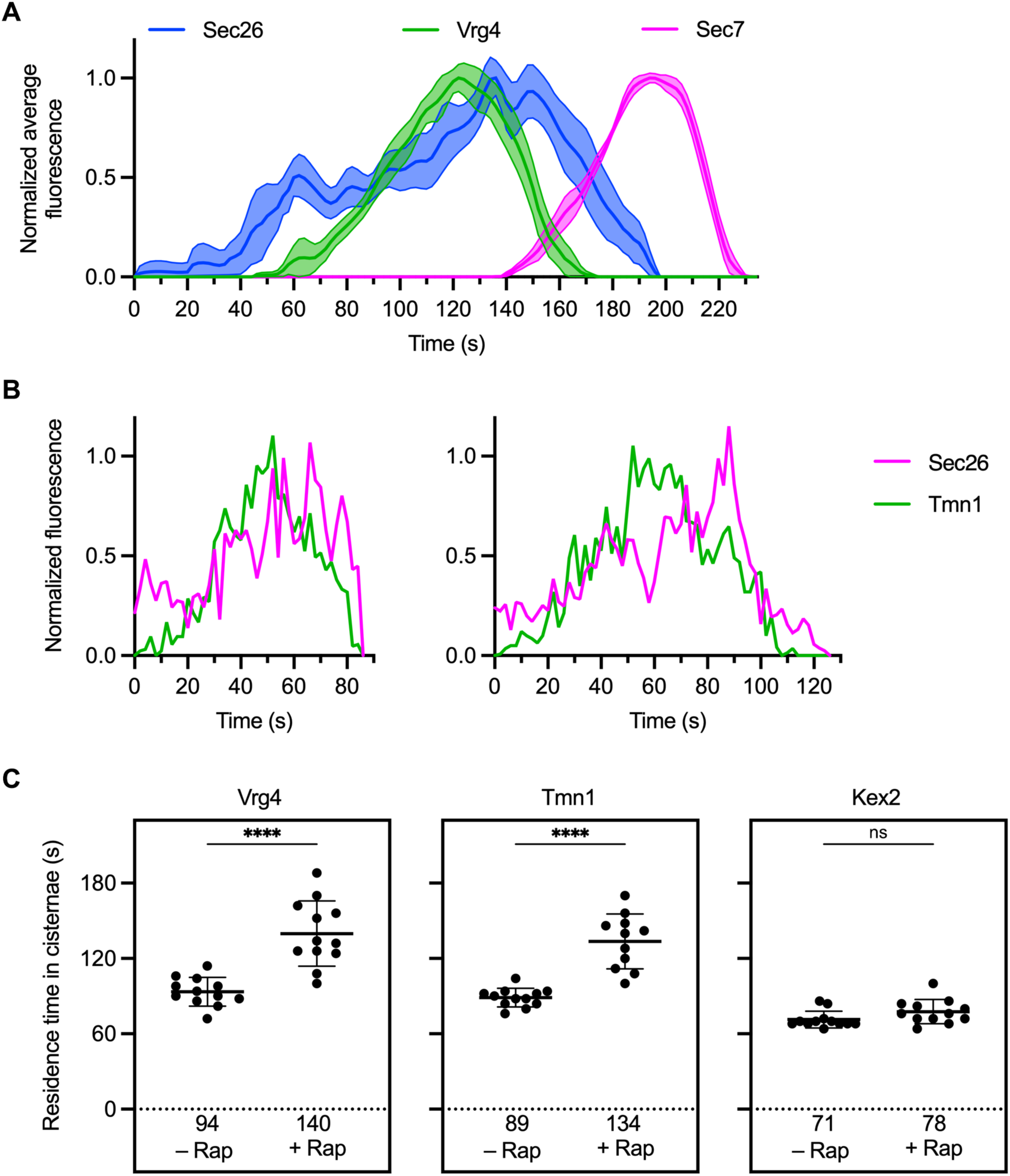
The intermediate recycling pathway depends on COPI. **(A)** Golgi maturation kinetics of the COPI subunit Sec26 relative to Vrg4 and Sec7. Shown are normalized and averaged traces for 11 individual cisternae marked with GFP-tagged Vrg4, mScarlet-tagged Sec7, and HaloTag-labeled Sec26. **(B)** Kinetic traces for two representative cisternae illustrating that COPI persists long enough to account for the departure of Tmn1. The traces show only the final part of the Sec26 residence times on the cisternae. **(C)** Residence times of Vrg4, Tmn1, and Kex2 in Golgi cisternae after partial inactivation of COPI. For the indicated GFP-tagged Golgi proteins, the residence times were defined as the intervals between the first and last detectable signals in individual maturing cisternae. Where indicated, partial inactivation of COPI was achieved by incubating with rapamycin (“Rap”) for 4-7 min before the first detectable signals of the GFP-tagged Golgi protein. The mean residence times are displayed as thick horizontal bars and are listed numerically below the plots, and the standard deviations are displayed as thin horizontal bars. ****, significant at P value <0.0001; ns, not significant.

To determine whether COPI mediates recycling of Tmn1, we modified an earlier approach in which COPI was acutely inactivated (Haruki et al., 2008; Papanikou et al., 2015). The COPI subunit Sec21 was tagged with FRB while a ribosomal subunit was tagged with FKBP, with the result that rapamycin treatment led to ribosomal association and functional inactivation of COPI. Maximal inactivation was previously seen after 10-15 min of rapamycin treatment, but by that time Golgi organization was disrupted (Papanikou et al., 2015). We therefore performed a partial inactivation of COPI by treating cells with rapamycin for only 4 min prior to kinetic mapping. The rationale was that we could track maturing cisternae in the usual way, except that reduced COPI function should slow the departure of proteins whose recycling is COPI-dependent. This approach was validated by examining the COPI-dependent protein Vrg4 and the COPI-independent protein Kex2 (Papanikou et al., 2015). In the absence of rapamycin, the average residence time of Vrg4 in maturing cisternae was 94 s, whereas in the presence of rapamycin, the average residence time of Vrg4 increased about 1.5-fold to 140 s (Fig. 6 C and Fig. S4, top and Video 4 and Video 5). By contrast, the average residence time of Kex2 was not significantly altered by rapamycin (Fig. 6 C and Fig. S4, bottom). This result indicates that partial inactivation of COPI selectively affected proteins that recycled in a COPI-dependent manner. For Tmn1, the average residence time was 89 s in the absence of rapamycin and 134 s in the presence of rapamycin (Fig. 6 C and Fig. S4, middle). As for Vrg4, the increase was about 1.5-fold. These results strongly suggest that COPI mediates the traffic of Tmn1 and other proteins that follow the intermediate recycling pathway.

### Vrg4-containing vesicles and intermediate pathway vesicles have different cargo compositions despite their shared dependency on COPI

The distinct kinetic signatures of Vrg4 and Tmn1 (see Fig. 4A) could reflect two separate COPI-dependent recycling pathways. If so, then we would expect to find additional resident Golgi transmembrane proteins that recycle together with Vrg4. A candidate was the mannosyltransferase Kre2 (also known as Mnt1; Lussier et al., 1995; Cho et al., 2000). Indeed, the kinetics of Kre2 arrival and departure were virtually identical to those of Vrg4 (Fig. 7 A, left). A second candidate was Pmr1, which also supports glycosylation, in this case by transporting divalent cations into the Golgi lumen (Antebi and Fink, 1992; Dürr et al., 1998). The kinetics of Pmr1 arrival and departure were very similar to those of Vrg4 (Fig. 7 A, right). We therefore predicted that Kre2 and Pmr1 would be found in the same vesicles as Vrg4.

**Figure 7.**
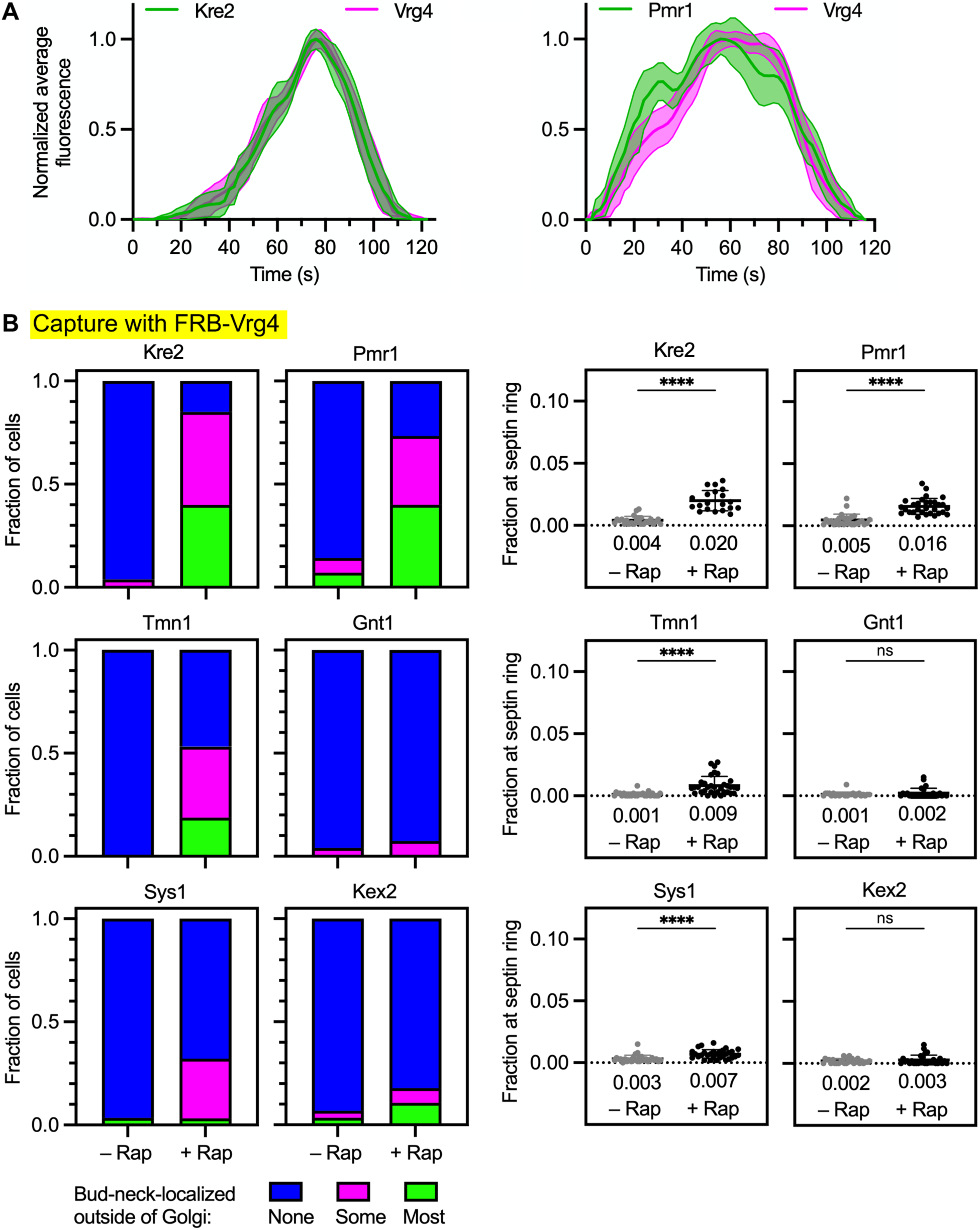
A subset of COPI cargoes can be specifically captured at the bud neck together with Vrg4. **(A)** Golgi maturation kinetics of HaloTag-labeled Vrg4 compared to GFP-tagged Kre2 or Pmr1. Shown are normalized and averaged traces for 12 individual cisternae for the Vrg4/Kre2 comparison and 11 individual cisternae for the Vrg4/Pmr1 comparison. The strain expressing tagged Kre2 carried the *vps10-104* mutation due to the luminal GFP tag. **(B)** Capture with FRB-Vrg4 of Kre2 and Pmr1, weak capture with FRB-Vrg4 of Tmn1 and Sys1, and no capture with FRB-Vrg4 of Gnt1 or Kex2. Categorical and numerical quantifications were performed as in Fig. 2 A. ****, significant at P value <0.0001; ns, not significant.

This prediction was tested by establishing a vesicle capture assay in which Vrg4 was tagged on its cytosolic N-terminus with FRB. To verify that Vrg4-containing vesicles could be captured, Vrg4 was tagged with both FRB and GFP. Upon rapamycin addition, the dual-tagged Vrg4 accumulated in vesicles at the bud neck as judged by fluorescence imaging (Fig. S5 A). Cryo-ET confirmed that capture of FRB-Vrg4 resulted in the appearance of vesicles at the bud neck (Fig. S5 B and Video 6). In cells expressing Vrg4 tagged with FRB alone, either Kre2, Pmr1, Tmn1, Gnt1, Sys1, or Kex2 was tagged with GFP. After rapamycin addition, only Kre2 and Pmr1 showed robust accumulation at the bud neck (Fig. 7 B and Fig. S5 C). No signal was seen with Gnt1 or Kex2, and weak signals were seen with Tmn1 and Sys1 (Fig. 7 B). Our interpretation is that Kre2 and Pmr1 frequently travel in vesicles together with Vrg4. The weak signals seen with Tmn1 and Sys1 might reflect occasional leakage of Vrg4 into intermediate pathway vesicles and/or occasional packaging of Tmn1 and Sys1 into early pathway vesicles. Taken together, the kinetic and vesicle capture data suggest that the cargoes recycled by the intermediate COPI-dependent pathway are at least partially distinct from those recycled by the early COPI-dependent pathway.

## Discussion

The polarized distribution of resident Golgi proteins confers temporal order on the events that occur during cisternal maturation, so a mechanistic understanding of Golgi polarity is crucial. We have proposed that Golgi polarity is generated by multiple membrane recycling pathways that operate with different kinetics during the life cycle of a cisterna (Pantazopoulou and Glick, 2019). Some recycling pathways act within the Golgi to deliver proteins from older to younger cisternae, and some recycling pathways act between the Golgi and other compartments (Fig. 8).

**Figure 8.**
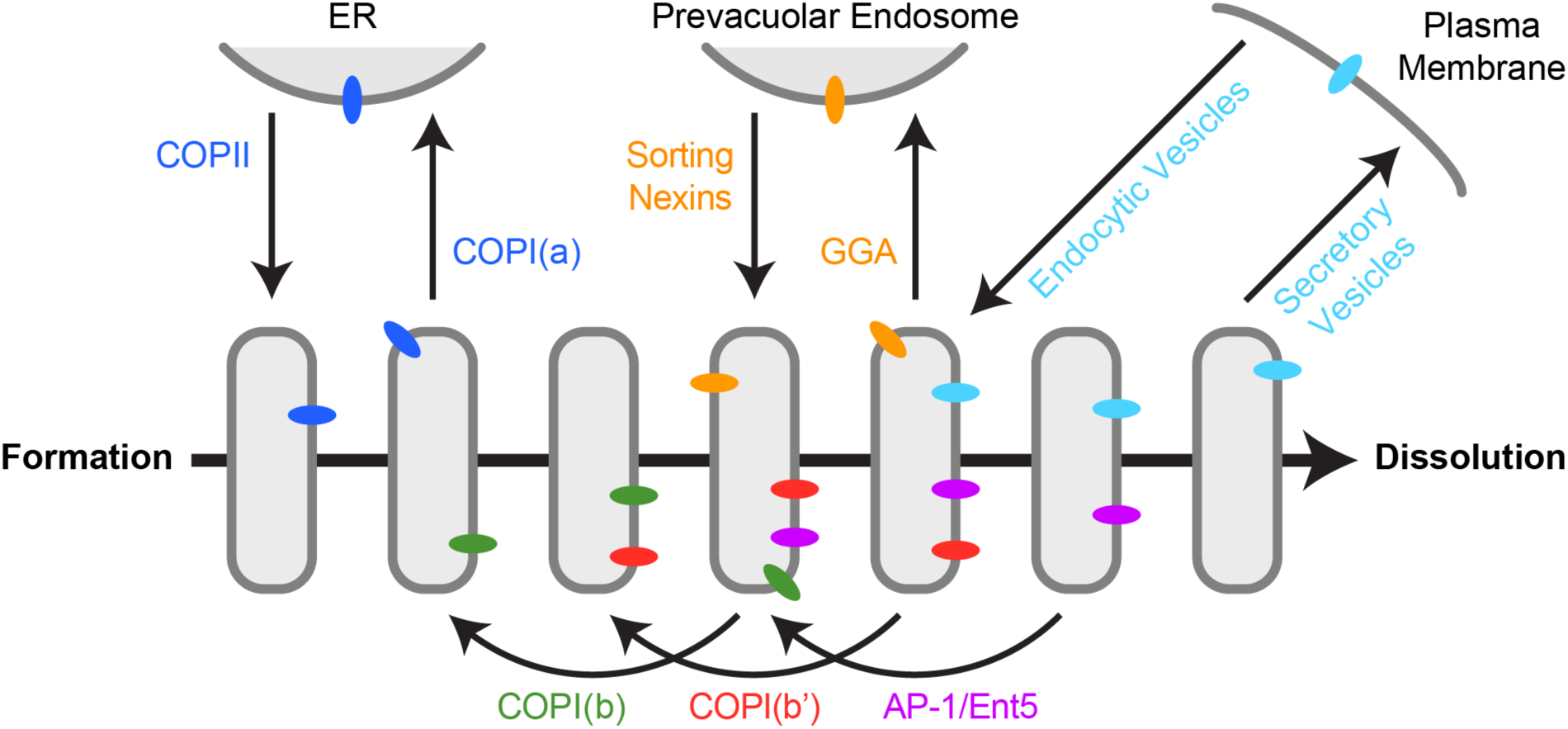
Summary of known recycling pathways that deliver membrane proteins to the yeast Golgi. The horizontal arrow depicts a timeline of cisternal maturation. During the lifetime of a cisterna, transmembrane proteins are delivered in six recycling pathways: (1) Some proteins (dark blue) are delivered to newly formed Golgi cisternae in COPII vesicles, and then those proteins recycle to the ER in COPI vesicles in a pathway that we term “COPI(a)”. (2) At an early stage of maturation, some resident Golgi proteins (green) recycle from older to younger cisternae in COPI vesicles in a pathway that we term “COPI(b)”. (3) At an intermediate stage of maturation, some resident Golgi proteins (red) recycle from older to younger cisternae in COPI vesicles in a pathway that we term “COPI(b’)”. (4) At an intermediate stage of maturation, some proteins (orange) are delivered to prevacuolar endosomes in clathrin-coated vesicles generated with the aid of the GGA adaptors, and then those proteins recycle to the Golgi in carriers generated with the aid of sorting nexins. (5) At a late stage of maturation, some resident Golgi proteins (magenta) recycle from older to younger cisternae in clathrin-coated vesicles generated with the aid of the AP-1 and Ent5 adaptors in a pathway that we term “AP-1/Ent5”. (6) At a terminal stage of maturation, some proteins (light blue) travel in secretory vesicles to the plasma membrane, and then those proteins recycle to the Golgi in endocytic vesicles.

What defines a Golgi recycling pathway? Our perspective is based on two assumptions. First, each recycling pathway is assumed to employ a specific traffic machinery that mediates the formation, tethering, and fusion of transport vesicles at a particular stage of maturation. Second, each recycling pathway is assumed to generate a distinct population of vesicles carrying a particular set of resident Golgi proteins. For example, our earlier work indicated that the COPI coat is involved in the recycling of Vrg4, which is present early in maturation, and that the clathrin adaptors AP-1 and Ent5 are involved in the recycling of Kex2, which is present late in maturation (Papanikou et al., 2015; Casler et al., 2022). A complication is that an individual component of the traffic machinery might function in multiple pathways. Examples are the role of COPI in both intra-Golgi and Golgi-to-ER traffic (Barlowe and Miller, 2013) and the ability of certain Golgi-associated tethering proteins to capture more than one class of transport vesicles (Wong and Munro, 2014). The likely implication is that a given recycling pathway harnesses a unique combination of traffic machinery components.

To characterize Golgi recycling pathways in yeast, we compare the kinetic signatures of various resident Golgi proteins, and we identify transmembrane proteins that can serve as reference markers for different pathways. This approach too has a complication: some resident Golgi proteins might follow more than one recycling pathway. The fidelity of partitioning into a given pathway is unlikely to be perfect, so many resident Golgi proteins probably follow a major recycling pathway as well as one or more minor salvage pathways (Glick, 2025). For example, a major pathway for recycling a set of Golgi proteins may also serve to salvage other Golgi proteins that recycle in an earlier pathway but occasionally leak downstream. Moreover, some resident Golgi proteins could partition at high frequency into more than one recycling pathway. This type of multi-pathway recycling might occur normally, e.g., if a resident Golgi protein functions at more than one stage of maturation. Alternatively, fluorescent tagging of a resident Golgi protein might perturb recycling in the major pathway, thereby pushing a fraction of the tagged molecules into a salvage pathway. Because of these caveats, the kinetic signatures of resident Golgi proteins are sometimes challenging to interpret, and additional types of data are needed.

We therefore developed a new functional test that is complementary to kinetic analysis. A specific population of Golgi-derived vesicles is captured at the yeast bud neck by chemically-induced dimerization (Fegan et al., 2010). For this purpose, an FRB-tagged Golgi protein in the vesicles heterodimerizes with an FKBP-tagged septin. Then fluorescence microscopy is used to determine which other resident Golgi proteins are co-captured in the same vesicles as the FRB-tagged protein. Table 1 lists the resident Golgi transmembrane proteins that were examined here, together with their inferred intra-Golgi recycling pathways as depicted in Fig. 8.

The vesicle capture assay gives meaningful results. For example, when Kex2 is tagged with FRB, there is efficient co-capture of Ste13 and Stv1, which have the same kinetic signature and the same AP-1/Ent5 dependency as Kex2 (Casler et al., 2022), and there is no co-capture of Vrg4, which has an earlier kinetic signature and a COPI dependency (Papanikou et al., 2015). But this assay also produces some ambiguous results as indicated below. We can envision possible sources of ambiguity. Most notably, if the FRB-tagged protein is occasionally missorted into a second type of vesicle, then that second type of vesicle will sometimes be captured at the bud neck, yielding a detectable signal. When interpreting the vesicle capture data, we compare different resident Golgi proteins in order to define stronger versus weaker co-capture, and we evaluate the findings in the context of kinetic analysis. This exercise has led to the following conclusions.

The available evidence points to a single intra-Golgi recycling pathway mediated by AP-1/Ent5. Our reference protein for this pathway is Kex2, which exhibits perturbed traffic in cells lacking AP-1 and Ent5 but not in cells with inactivated COPI (Papanikou et al., 2015; Casler et al., 2022). We previously suggested that a second AP-1/Ent5-dependent pathway might recycle Sys1 upstream of Kex2 (Casler et al., 2022). The reasoning was that (a) Sys1 arrived at maturing cisternae earlier than Kex2, and (b) Sys1 traffic was altered in cells lacking AP-1 and Ent5. However, the vesicle capture assay reveals that Sys1 is co-captured with Kex2, leading us to suspect that Sys1 is not a suitable marker for a unique recycling pathway. The combined kinetic and functional data support the idea that Sys1 partitions between two recycling pathways: a late pathway marked by Kex2, and an upstream pathway with intermediate kinetics. This dual partitioning could reflect the physiological behavior of Sys1, but it might be due to the cytosolic tag. When cytosolically-tagged Aur1 (Casler et al., 2022) is analyzed by kinetic mapping and by the vesicle capture assay, it shows the same apparent dual partitioning as Sys1. A possible interpretation is that Sys1 and Aur1 normally recycle in the intermediate pathway and that entry into this pathway is sensitive to bulky cytosolic tags.

The dual partitioning of Sys1 and Aur1 prompted us to look for other resident Golgi proteins that recycle mainly in the intermediate pathway. We identified Tmn1, which can be luminally tagged. Tmn1 arrives at maturing cisternae before Kex2, and Tmn1 departs before Kex2. The relative kinetic signatures of Tmn1, Sys1, and Kex2 suggest that tagged Sys1 partitions between an intermediate pathway marked by Tmn1 and a late pathway marked by Kex2. This scenario is bolstered by the finding that Tmn1 is co-captured strongly in vesicles containing FRB-tagged Sys1 but only very weakly in vesicles containing FRB-tagged Kex2. Gnt1 can also be luminally tagged, and it resembles Tmn1 with regard to its kinetic behavior and its selective co-capture with FRB-tagged Sys1. Based on these data, we propose the existence of an intermediate recycling pathway for which Tmn1 can serve as the reference marker.

Is the intermediate pathway mediated by COPI, or is it mediated by AP-1/Ent5? Kinetic analysis indicated that COPI is present on a maturing cisterna throughout the Tmn1 departure phase whereas AP-1 is absent from a maturing cisterna until the very end of the Tmn1 departure phase. This result makes COPI a much better candidate than AP-1/Ent5 for generating vesicles of the intermediate pathway. One way to test this prediction is to block endocytosis in cells lacking AP-1 and Ent5. In such mutant cells, proteins that normally recycle in the AP-1/Ent5 pathway apparently travel to the plasma membrane and then return to the Golgi in endocytic vesicles (Day et al., 2018; Casler et al., 2022). We found that blocking endocytosis in cells lacking AP-1 and Ent5 strongly perturbs the traffic of Kex2 and Ste13 but not of Vrg4 or Tmn1. A moderate effect is seen with Sys1, consistent with the interpretation that some of the tagged Sys1 molecules normally recycle in the AP-1/Ent5 pathway. A second way to test which type of coat generates vesicles of the intermediate pathway is to reduce the activity of COPI by anchoring it to ribosomes (Papanikou et al., 2015). Partial inactivation of COPI slows the departure of Vrg4 and Tmn1 from maturing cisternae but has no effect on the departure of Kex2. That result fits with reports that Vrg4 and Tmn1 have COPI recognition signals (Abe et al., 2004; Woo et al., 2015). The combined data indicate that Tmn1, and presumably also Gnt1, follow a COPI-dependent recycling pathway with intermediate kinetics.

Because both Vrg4 and Tmn1 show COPI dependency but they arrive at a maturing cisterna at different times, those two proteins likely recycle in distinct COPI-mediated pathways. To test this prediction, we performed vesicle capture assays using FRB-tagged Vrg4. Kre2 and Pmr1 have kinetic signatures similar to that of Vrg4, and they show relatively strong co-capture with Vrg4, suggesting that they are present in the same vesicles as Vrg4. By contrast, Gnt1 and Kex2 show no co-capture with Vrg4, consistent with the idea that the intermediate and late recycling pathways are distinct from the early pathway marked by Vrg4. Relatively weak capture with Vrg4 was seen for Tmn1 and Sys1. This result might reflect occasional leakage of Vrg4 into the intermediate pathway or occasional packaging of Tmn1 and Sys1 into vesicles of the early pathway. Thus, while some of the individual results are not definitive, the combination of kinetic and functional analyses supports the existence of two COPI-dependent intra-Golgi pathways marked by Vrg4 and Tmn1. One could envision that Vrg4 follows a Golgi-to-ER pathway while Tmn1 follows an intra-Golgi pathway. But if Vrg4 recycled through the ER, it should be visible in the youngest Golgi cisternae, and that result is not observed (Casler et al., 2019). Instead, the appearance of Vrg4 is preceded by that of certain other Golgi markers (Tojima et al., 2024)—including COPI itself, as shown here—suggesting that Vrg4 recycles within the Golgi. Thus, Vrg4 and Tmn1 apparently recycle in successive intra-Golgi pathways, which we provisionally term COPI(b) and COPI(b’), respectively (Fig. 8 and Table 1).

Interestingly, Golgi proteins that seem to follow the same recycling pathway, as judged by matching arrival kinetics and by co-capture assays, do not always show matching departure kinetics. For example, in the COPI(b’) pathway, Gnt1 arrives at the same time as Tmn1 but typically departs slightly before Tmn1. This finding can be rationalized by assuming that Gnt1 and Tmn1 compete for packaging into outgoing COPI(b’) vesicles, with Gnt1 being the stronger competitor and therefore departing sooner (Glick et al., 1997). In this scenario, COPI(b’) vesicles might be somewhat heterogeneous in composition but they would all fuse with a cisterna at the same stage of maturation. The phenomenon of variable departure kinetics will be explored more fully in future work (C. Lee-Smith and A. Krahn, in preparation).

Golgi proteins that follow the same recycling pathway(s) would be expected to arrive with similar kinetics. For the most part, that pattern seems to hold true, with vesicle capture assays supporting the hypothesis that proteins arriving at the same time are indeed in the same vesicles. An apparent exception is Sys1 and Tmn1 in the COPI(b’) pathway, because accumulation of Sys1 lags behind that of Tmn1 even though Tmn1 can be co-captured with Sys1. However, Sys1 is unusual because it partitions between the COPI(b’) and AP-1/Ent5 pathways. Our analysis method may be subtly misleading in this case, based on the following logic. If the partitioning varies such that for some cisternae, a large fraction of the Sys1 molecules follow the COPI(b’) pathway together with Tmn1, whereas for other cisternae, only a small fraction of the Sys1 molecules follow the COPI(b’) pathway while the rest follow the downstream AP-1/Ent5 pathway, then the averaged kinetic signature will show Sys1 lagging behind Tmn1. This explanation is consistent with the observed variability in the individual traces comparing Sys1 with Tmn1. Another possible explanation is that Sys1 but not Tmn1 often departs from a cisterna immediately after arrival, with the result that Sys1 accumulates more slowly than Tmn1. In any case, further work is needed to gain a full understanding of Sys1 recycling.

The high-level summary of our findings is that at least two COPI-dependent pathways mediate intra-Golgi recycling early in maturation while an AP-1/Ent5-dependent pathway mediates intra-Golgi recycling late in maturation. It is interesting that even though yeast cells contain just a single species of the COPI coatomer (Gaynor et al., 1998), COPI mediates multiple pathways, including intra-Golgi recycling as well as Golgi-to-ER recycling. The implication is that COPI acts together with partner proteins to ensure specificity for both cargo capture and vesicle targeting. An example of a COPI partner protein is yeast Vps74, which is related to mammalian GOLPH3 and GOLPH3L (Schmitz et al., 2008; Tu et al., 2008; Welch et al., 2021). Vps74 and its mammalian orthologs interact with COPI and with a subset of the cargo proteins that are packaged into COPI vesicles. We speculate that additional COPI partner proteins assist in specifying the various COPI-dependent recycling pathways.

The insights described here bring us closer to a complete picture of the membrane recycling pathways at the yeast Golgi. A future goal is to identify the tethers and SNAREs that operate in each pathway. The resulting data set will be the framework for an integrated model of the Golgi system.

## Materials and methods

### Yeast growth and transformation

All strains used in this study were derivatives of JK9-3da (*leu2-3,112 ura3-52 rme1 trp1 his4*; Kunz et al., 1993) with the *pdr1Δ pdr3Δ* mutations to facilitate HaloTag labeling (Barrero et al., 2016; Casler et al., 2022). Rapamycin-resistant strains carrying the *fpr1Δ* and *TOR1-1* mutations were constructed as previously described (Papanikou et al., 2015). If a Golgi protein was luminally tagged with GFP or HaloTag, the strain carried the *vps10-104* mutation to prevent missorting to the vacuole (Fitzgerald and Glick, 2014; C. Lee-Smith, in preparation). Prior to experimental analysis, yeast cells were grown in nonfluorescent minimal glucose medium (NSD) (Bevis et al., 2002) in baffled flasks at 23°C with shaking at 200 rpm unless otherwise noted.

Yeast genes were tagged by chromosomal gene replacement using the pop-in/pop-out method (Rothstein, 1991; Rossanese et al., 1999). The plasmids used for those manipulations were constructed with the aid of SnapGene software (Dotmatics) and the Saccharomyces Genome Database (Wong et al., 2023), and they have been archived at Addgene together with annotated maps. Tagged Tmn1 was likely cleaved by Kex2 (Schimmöller et al., 1998), but the two fragments of the protein are predicted to remain associated based on an AlphaFold Server simulation (Abramson et al., 2024).

### Reagents

HaloTag ligands of JFX_646_ and JFX_650_ (Grimm et al., 2021), generously provided by Luke Lavis (Janelia Research Campus, Ashburn, VA), were dissolved in anhydrous DMSO (Invitrogen, catalog no. D12345) at a concentration of 1 mM, and single-use aliquots were stored at -80°C. These stock solutions were diluted 1000-fold to label cells at a final dye concentration of 1 µM. Rapamycin (LC Laboratories, catalog no. R-5000) was prepared as a 1 mg/ml solution in 90% ethanol, 10% Tween 20, and single-use aliquots were stored at -20°C. This stock solution was diluted 100-fold to treat cells at a final rapamycin concentration of 10 µg/ml. CK-666 (Sigma, catalog no. SML0006) was dissolved in anhydrous DMSO to a concentration of 50 mM, and single-use aliquots were stored at 4°C. This stock solution was diluted 500-fold to treat cells at a final CK-666 concentration of 100 µM.

### HaloTag labeling

Labeling of HaloTag constructs (Casler et al., 2022) was performed as follows. HaloTag ligand of JFX_646_ or JFX_650_ was diluted by adding 1 µl of a 1 mM DMSO stock solution to 300 µl of NSD. The resulting solution was cleared of precipitate by spinning at 17,000xg (13,000 rpm) in a microcentrifuge for 5 min. Then the supernatant was added to 700 µl of mid-log phase yeast culture to give a final concentration of 1 µM dye, and the cells were incubated for 30-60 min at 23°C with shaking. For live cell imaging experiments, excess dye was removed by filtration and washing with a 0.22-µm syringe filter (Millipore; catalog no. SLGV004SL). The washed cells on the filter were resuspended in NSD and adhered to a coverglass-bottom dish coated with concanavalin A (ConA) (Johnson and Glick, 2019). Confocal videos were then immediately acquired as described below.

### Live cell fluorescence microscopy and analysis

For 4D confocal microscopy, yeast cells were attached to ConA-coated coverglass-bottom dishes that were filled with NSD (Johnson and Glick, 2019). Imaging was performed at 23°C using a Leica Stellaris confocal microscope equipped with a 1.4-NA/63x oil objective. Imaging of cell volumes was performed with 40-80 nm pixels, a 0.30 µm z-step interval, and 20-30 optical sections. z-stacks were obtained at intervals of 2 s.

Confocal movies of Golgi cisternal maturation events were deconvolved using Huygens software (SVI) and quantified with custom ImageJ plugins as previously described (Johnson and Glick, 2019), except that Huygens software was employed for time-based correction of photobleaching prior to deconvolution. Normalization and averaging of kinetic traces from maturing Golgi cisternae were accomplished using custom ImageJ plugins (Casler et al., 2022).

### Vesicle capture assay

Cells grown overnight to an OD_600_ of ∼0.5 were stained with 1 µM JFX_650_ HaloTag ligand as described above, except that the final volume was 500 µl. After growth with shaking for 1 h at 23°C, either a 100-fold dilution of 90% ethanol, 10% Tween 20 was added as a control or rapamycin was added to a final concentration of 10 µg/ml, and the cells were incubated with shaking for an additional 5 min. Then fixation was performed by adding 250 µl of the culture while vortexing to 750 µl of 1.33-fold concentrated fixative to give final concentrations of 1% paraformaldehyde plus 0.1% glutaraldehyde (Electron Microscopy Sciences, diluted from freshly opened vials), 1 mM MgCl_2_, 50 mM potassium phosphate, pH 6.5. After fixation on ice for 1 h, the cells were washed twice by centrifuging for 2 min at 1500xg (4000 rpm) and resuspending in 500 µl of phosphate-buffered saline (PBS). Finally, the cells were centrifuged once again and resuspended in 20 µl PBS. Within 30 h of fixation, the fixed cells were compressed under coverslips and imaged with a Leica Stellaris confocal microscope using a 1.4-NA/63x oil objective with 40 nm pixels, a 0.20 µm z-step interval, and 21 optical sections. Cells were chosen for analysis if they were a mother-daughter cell pair with a joined septin ring, and if Golgi cisternae were largely absent from the bud neck region as determined by viewing the Golgi marker channel. The cell images were deconvolved using Huygens software (Johnson and Glick, 2019). A set of 20-30 cells was then analyzed by the two quantification methods described below.

For categorical quantification of vesicle capture, ImageJ was first used to process the deconvolved images of untreated and rapamycin-treated cells, as follows. The images were average projected, and the contrast of each fluorescence channel was enhanced by choosing a pixel saturation percentage: 0.3% for the red channel, 0.5% for the green channel, or 0.4% for the blue channel. Then the channels were merged to generate composite images. To ensure objective quantification, the Blind Analysis Tools (v1.0) of ImageJ were employed to hide image identities. Image names were encrypted using basic mode of the File Name Encrypter, and the Analyse & Decide tool was used to assign each budding cell by visual criteria to one of the following categories: 1) No bud-neck localized cargo outside of cisternae, 2) Minority of bud-neck localized cargo outside of cisternae, or 3) Majority of bud-neck localized cargo outside of cisternae.

For numerical quantification of vesicle capture, a confocal image stack of a cell was processed with a custom ImageJ plugin termed “Quantify Overlap”, available with source code from https://github.com/bsglicker/4D-Image-Analysis. With this plugin, the user chooses a region of interest (ROI), and the ROI measurements are summed for each image in the z-stack. As used here, the plugin creates a mask from the signal in the red channel and then compares the green signal within the mask to the total green signal in the ROI. The red channel mask can be modified by creating a second mask from the signal in the blue channel and then subtracting the blue channel mask from the red channel mask. Threshold levels are chosen empirically by the user for the red and blue channel masks, from a set of threshold options generated by a custom algorithm. For our purposes, the red threshold was set to the “Lower” level, and the blue threshold was set to the lowest “Basement” level to create an extensive mask that included all visible Golgi signal. A cell was used for further analysis only if subtraction of the blue channel mask removed less than 65% of the area from the red channel mask.

Representative fluorescence images chosen for display in figures were scaled to the full RGB dynamic range, and then the pixel values were multiplied by 1.5 to ensure adequate visibility of both bright and faint structures.

### AP-1/Ent5 dependency assay

To assess whether traffic of a Golgi protein was affected by loss of AP-1 and Ent5, 5-ml cultures of wild-type (*APL4 ENT5*) and mutant (*apl4Δ ent5Δ*) yeast strains expressing the GFP-tagged Golgi protein were grown overnight to an OD_600_ of ∼0.5. Two 500-µl samples were placed in culture tubes. One sample received 1 µl of DMSO as a control, and the other sample received 1 µl of 50 mM CK-666. After 5-6 min, 250 µl of each sample were transferred to ConA-coated coverglass-bottom dishes. After an additional 5 min, the medium was removed, and the cells were washed and then overlaid with NSD either lacking or containing CK-666 as appropriate. Confocal z-stacks were captured between 15-25 min after drug addition.

The z-stacks were deconvolved and average projected. For a given Golgi protein, the subsequent quantification was blinded with regard to control versus CK-666-treated samples using ImageJ as described above. Individual cells were visually examined and assigned to categories based on whether (a) most of the GFP-tagged protein was in Golgi-like puncta distributed throughout the cytoplasm with no visible signal at secretion sites, or (b) most of the GFP-tagged protein was concentrated at secretion sites (i.e., sites of polarized growth) with little signal in Golgi-like puncta elsewhere in the cytoplasm, or (c) the GFP-tagged protein was distributed in a hybrid pattern between Golgi-like puncta and secretion sites.

In control experiments, the ability of CK-666 to inhibit endocytosis was confirmed by showing that this drug blocked internalization of the dye FM 4-64 (Casler et al., 2022).

### COPI inactivation assay

Live yeast cells expressing Sec21-FRB and Rpl13A-FKBPx2 (Papanikou et al., 2015) plus mScarlet-tagged Sec7 and a GFP-tagged Golgi protein were prepared for confocal imaging by adhering them to ConA-coated coverglass-bottom dishes (Johnson and Glick, 2019). To inhibit COPI activity by anchoring to ribosomes, the medium was replaced with NSD containing 10 µg/ml rapamycin. For the control sample, the medium was replaced with NSD containing 0.9% ethanol, 0.1% Tween 20. Cells were imaged for 6 min starting 4 min after rapamycin addition. Maturation events beginning between 4 and 7 min after rapamycin addition were analyzed to determine the cisternal residence times of the GFP-tagged Golgi protein, where the cisternal residence time for a given cisterna was defined as the interval between the first and last visible GFP signals.

### Cryo-electron tomography

Quantifoil R2/2 grids on 200 copper mesh were prepared by glow discharging them. Log-phase yeast cultures in NSD, either mock-treated or treated for 5 min with 10 µg/ml rapamycin, were applied to the grids, which were then cryopreserved using a Vitrobot Mark IV (Thermo Fisher). The sample chamber was kept at 16 °C and 100% humidity with a 60 s wait time, and grids were blotted for 11 s before being plunged into liquid ethane. Grids were stored under liquid nitrogen.

For cryo-FIB milling, grids were clipped in AutoGrid Rings (Thermo Fisher) with a cut-out to allow for shallower milling angles. Milling was performed on an Aquilos 2 dual-beam instrument (Thermo Fisher). Samples were sputter-coated in-column with platinum for 20 s at 10 Pa and 20 mA, and then they were coated with a layer of organometallic platinum for 35 s using the gas injection system within the instrument. Targets were identified and milled using Maps software and milled using AutoTEM software (Thermo Fisher). The automated protocol included milling of micro-expansion joints at 0.5 nA, followed by rough, fine, and very fine milling at 0.5, 0.3, and 0.1 nA, respectively, followed by two polishing steps at 50 and 30 pA. The result was a lamella with a target thickness of 200-220 nm. Milled samples were stored under liquid nitrogen.

For cryo-ET and modeling, data were collected on a Titan Krios G3i at 300 kV (Thermo Fisher) with a Gatan K3 direct detection camera in CDS mode with the initial dose rate target on the detector between 7.5-8.5 electrons per pixel per second and with the Gatan BioQuantum-K3 energy slit width set to 20 eV. Tomography 5 software (Thermo Fisher) was used for collection. The tilt series was acquired with 3° steps in a bidirectional collection scheme (Hagen et al., 2017) beginning with a lamella pre-tilt of ±9° and extending to ±54°. The total dose for each tilt series was 120 e-/Å^2^. Tilts were acquired with a pixel size of 4.45 Å and with 5 µm of defocus. Tilt series data were initially reconstructed using Tomo Live software (Comet et al., 2024; Thermo Fisher), which allowed for on-the-fly motion correction, alignment, and reconstruction of tomograms using the Simultaneous Iterative Reconstruction Technique (SIRT; Gilbert, 1972). Target features were identified in the tomograms generated by Tomo Live, and those features were segmented using the 3dmod package in IMOD (Kremer et al., 1996). At least five lamellae from large budded cells were examined for each experimental condition.

To estimate the maximal distance between the plasma membrane beneath the septin ring and a captured vesicle, we used AlphaFold 3 (https://alphafoldserver.com) to simulate the structures of FKBP-tagged Shs1 and the FRB-tagged cytosolic portions of Kex2, Sys1, and Vrg4. Assuming that all of the flexible protein regions were maximally extended, the estimated lengths were 48 nm for Shs1-FKBP, 58 nm for Kex2-FRB, 35 nm for Sys1-FRB, and 18 nm for FRB-Vrg4.

### Statistical analysis

For vesicle capture experiments, Microsoft Excel was used to compile the quantification data. Numerical quantification of mean percentages was performed by first excluding values from cells that exhibited high levels of cisternal interference at the bud neck, where a high level was defined as removal of at least 65% of the red septin mask by subtraction of the blue Golgi mask. The data were then transferred to Prism software (Dotmatics) for plotting and for computing standard deviations. For cisternal residence time measurements, the data were directly transferred to Prism. The P values of scatter plots were calculated in Prism using Welch’s t test.

## Acknowledgments

This work in the B.S.G. laboratory was supported by NIH grant R35 GM144050. A.H.K. was supported by NIH training grant T32 GM007183. Alec Buttner assisted with some of the initial vesicle capture experiments that revealed unexpected complexity with Sys1 trafficking. Thanks for assistance with fluorescence microscopy to Vytas Bindokas, Christine Labno, and Lorraine Horwitz at the Integrated Light Microscopy Core Facility (RRID:SCR_019197), which is supported by the NIH-funded Cancer Center Support Grant P30 CA014599. Thanks for assistance with electron microscopy to Yimei Chen, Tera Lavoie, and Joshua Fisher at the Advanced Electron Microscopy Core Facility (RRID:SCR_019198). We thank Kasey Day for generating the Sys1-FRB construct and Luke Lavis for providing the JFX_650_ dye. The authors declare no competing financial interests.

## Author Contributions

Adam Krahn: conceptualization, data curation, formal analysis, investigation, methodology, validation, visualization, and writing—original draft, review, and editing. Areti Pantazopoulou: conceptualization, data curation, formal analysis, investigation, methodology, validation, visualization, and writing—review and editing. Jotham Austin II: conceptualization, data curation, formal analysis, investigation, methodology, validation, visualization, and writing—review and editing. Natalie Johnson: conceptualization, data curation, formal analysis, investigation, methodology, validation, visualization, and writing—review and editing. Conor Lee-Smith: conceptualization, data curation, formal analysis, investigation, methodology, validation, visualization, and writing—review and editing. Benjamin Glick: conceptualization, formal analysis, funding acquisition, project administration, resources, supervision, validation, and writing—original draft, review, and editing.

## Abbreviations

ConA: concanavalin A
cryo-ET: cryo-electron tomography
FKBP: FK506-binding protein
FRB: FK506-rapamycin binding domain
NSD: nonfluorescent minimal glucose medium
SIRT: Simultaneous Iterative Reconstruction Technique;
TGN: *trans*-Golgi network

---

**Figure S1.**
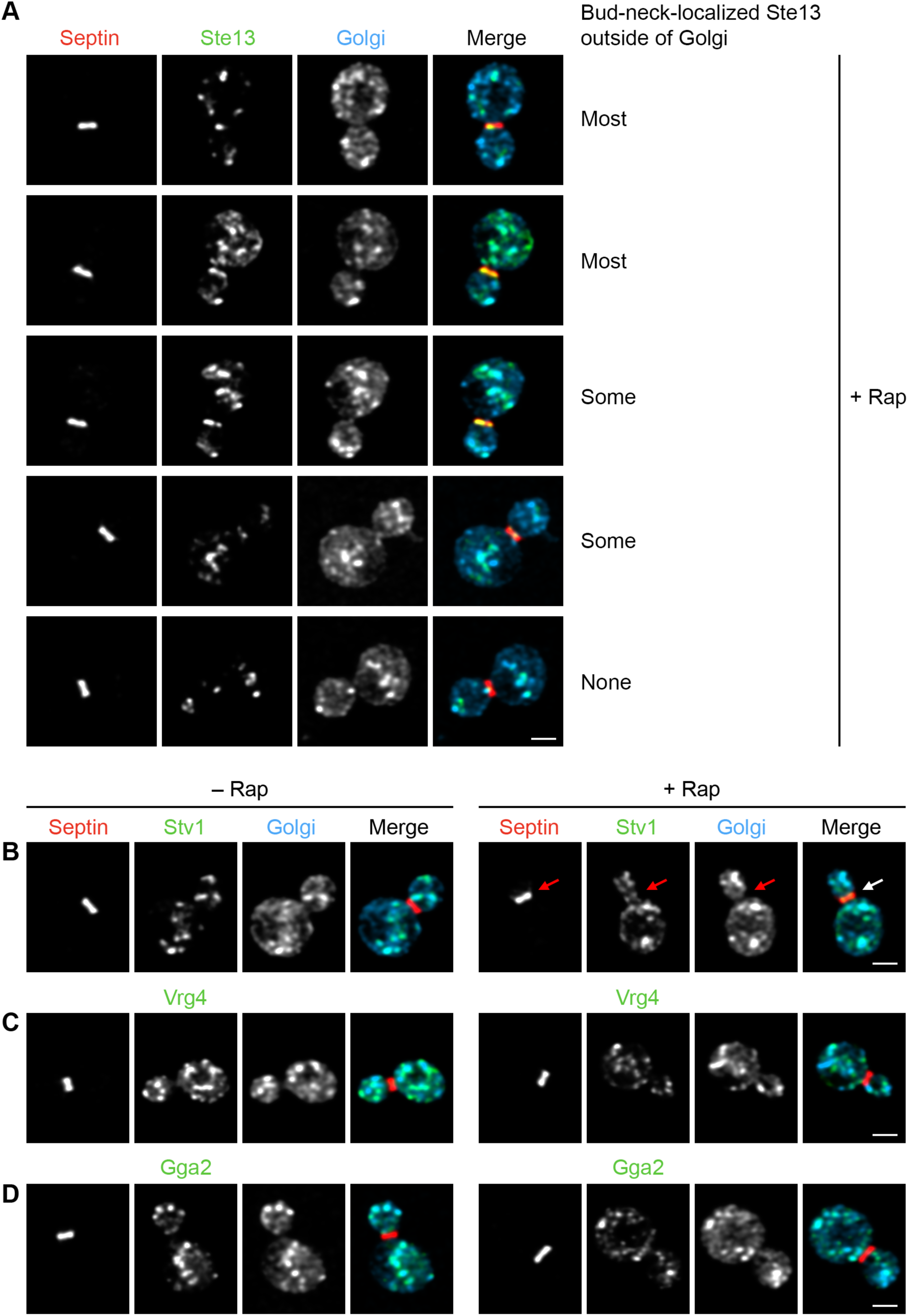
Supplementary data for Fig. 2. In each part of the figure, Golgi cisternae were marked by HaloTag-labeled Ric1 and Sec7 (blue), and the scale bar is 2 µm. **(A)** A panel of representative images showing capture with Kex2-FRB of GFP-tagged Ste13 (green) by an FKBP-tagged septin (red) after treatment for 5 min with rapamycin (“Rap”). The labels “Most”, “Some”, and “None” mark representative examples for the image categories used in Fig. 2 A. **(B)** Capture with Kex2-FRB of GFP-tagged Stv1 (green) by an FKBP-tagged septin (red) after treatment for 5 min with rapamycin. Arrows indicate non-Golgi signal at the bud neck. **(C)** No capture with Kex2-FRB of GFP-tagged Vrg4 (green) by an FKBP-tagged septin (red) after treatment for 5 min with rapamycin. **(D)** Minimal capture with Kex2-FRB of GFP-tagged Gga2 (green) by an FKBP-tagged septin (red) after treatment for 5 min with rapamycin.

**Figure S2.**
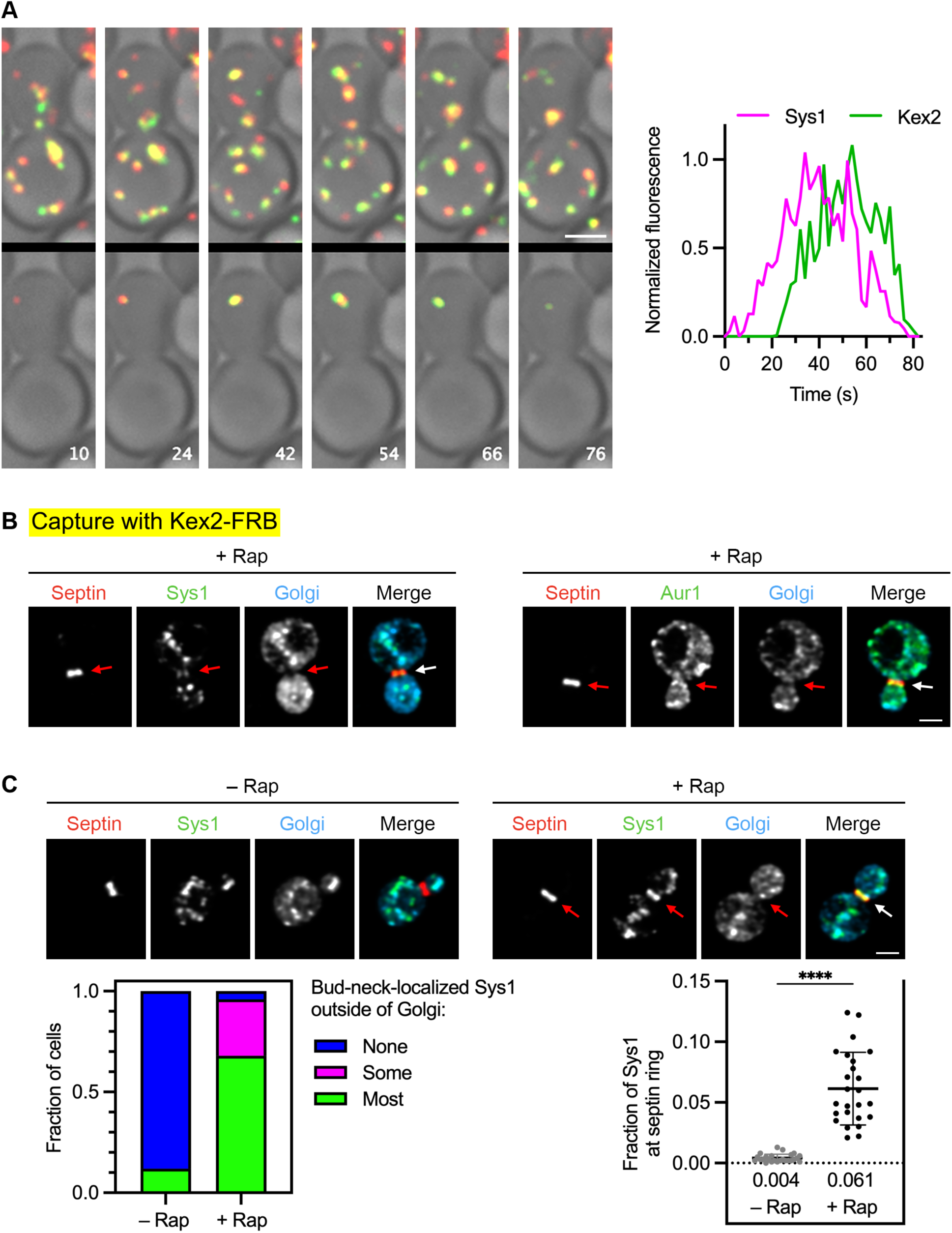
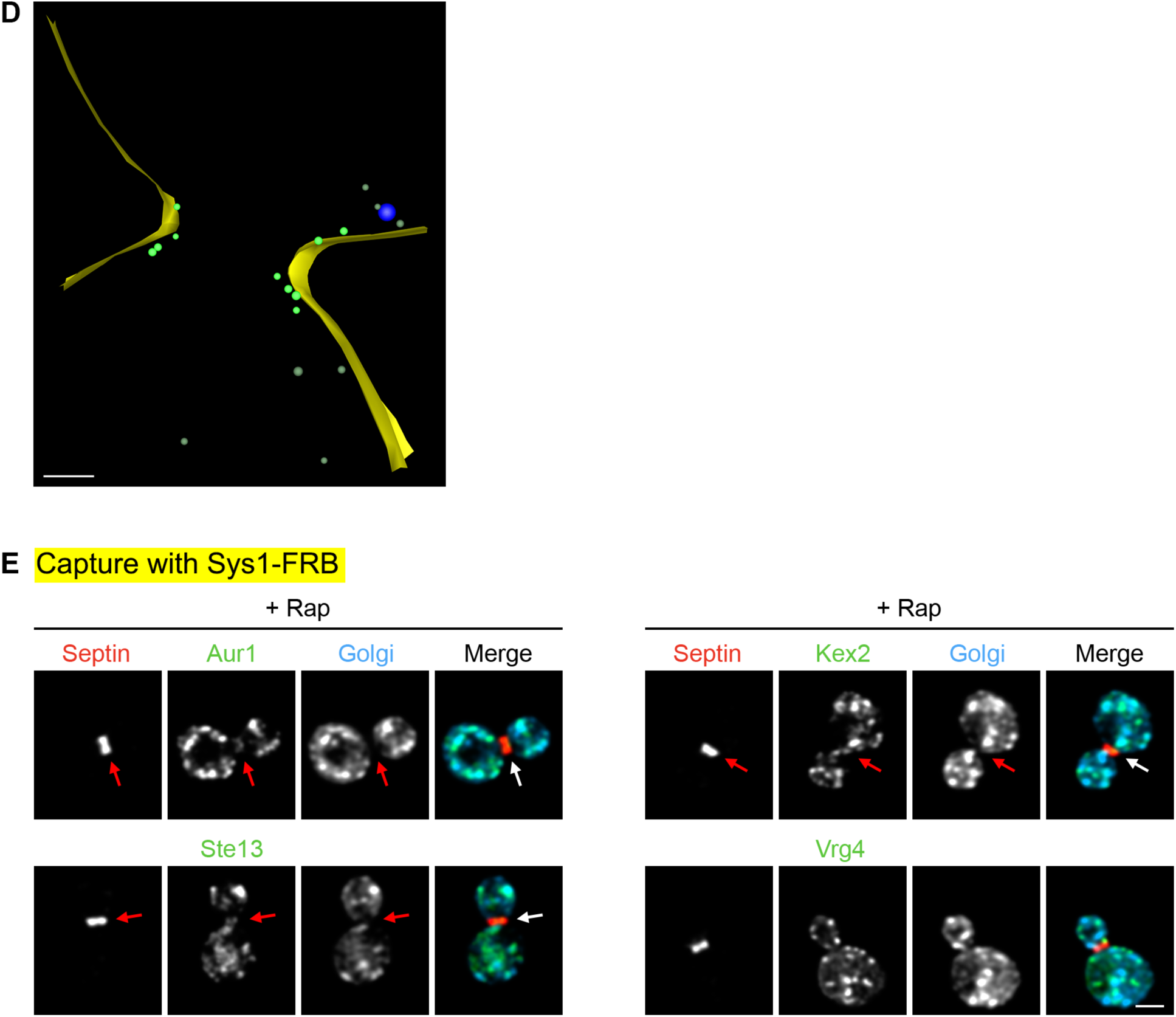
Supplementary data for Fig. 3. **(A)** Frames from a representative 4D confocal movie of Sys1-HaloTag (red) and Kex2-GFP (green), and kinetic traces from an individual cisterna in the movie. Depicted at the left are average projected z-stacks at the indicated time points from Video 2. The upper row shows the complete projections, and the lower row shows edited projections that include only the cisterna that was tracked. Scale bar, 2 µm. Plotted at the right are normalized fluorescence intensities for the cisterna tracked in the movie. **(B)** Representative images showing capture with Kex2-FRB of GFP-tagged Sys1 or Aur1 (green) by an FKBP-tagged septin (red) after treatment for 5 min with rapamycin. Arrows indicate non-Golgi signal at the bud neck. **(C)** Rapamycin-dependent capture by an FKBP-tagged septin (red) of Sys1-FRB-GFP (green). Arrows indicate non-Golgi signal at the bud neck after treatment for 5 min with rapamycin. At the lower left, capture of Sys1-FRB-GFP at the bud neck was quantified by assigning cells to categories as described in Fig. 1 C. At the lower right, capture of Sys1-FRB-GFP at the bud neck was quantified numerically as described in Fig. 1 C. ****, significant at P value <0.0001. **(D)** Cryo-ET of vesicles captured at the bud neck using Sys1-FRB. A log-phase culture of cells expressing Sys1-FRB and Shs1-FKBP was treated with rapamycin for 5 min followed by cryopreservation and processing for cryo-ET. Shown is the model of a SIRT-reconstructed tomogram from a large budded cell. The full data set is shown in Video 3. A vesicle was counted as putatively captured if its membrane was no more than 83 nm from a point on the plasma membrane within 200 nm from the center of the bud neck. For the 4 non-rapamycin-treated cells examined, the number of vesicles meeting this criterion ranged from 0 to 2 (mean = 1.0). For the 4 rapamycin-treated cells examined, the number of putatively captured vesicles ranged from 6 to 13 (mean = 10.0). Scale bar, 250 nm. See Fig. 1 D for further details. **(E)** Representative images showing capture with Sys1-FRB of GFP-tagged Aur1, Kex2, or Ste13 (green) by an FKBP-tagged septin (red), and minimal capture with Sys1-FRB of Vrg4 (green) by an FKBP-tagged septin (red), all after treatment for 5 min with rapamycin. Arrows indicate non-Golgi signal at the bud neck.

**Figure S3.**
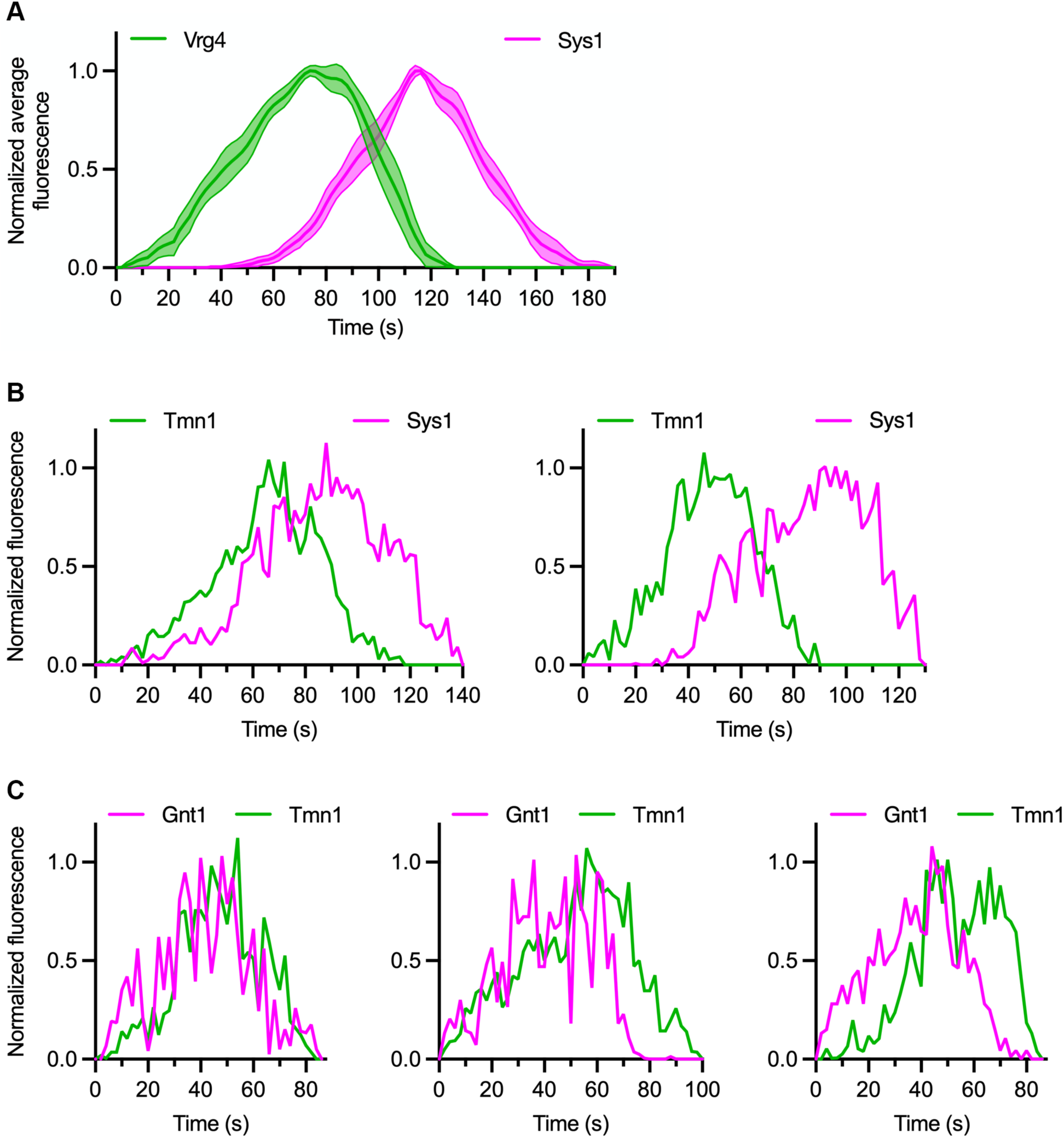
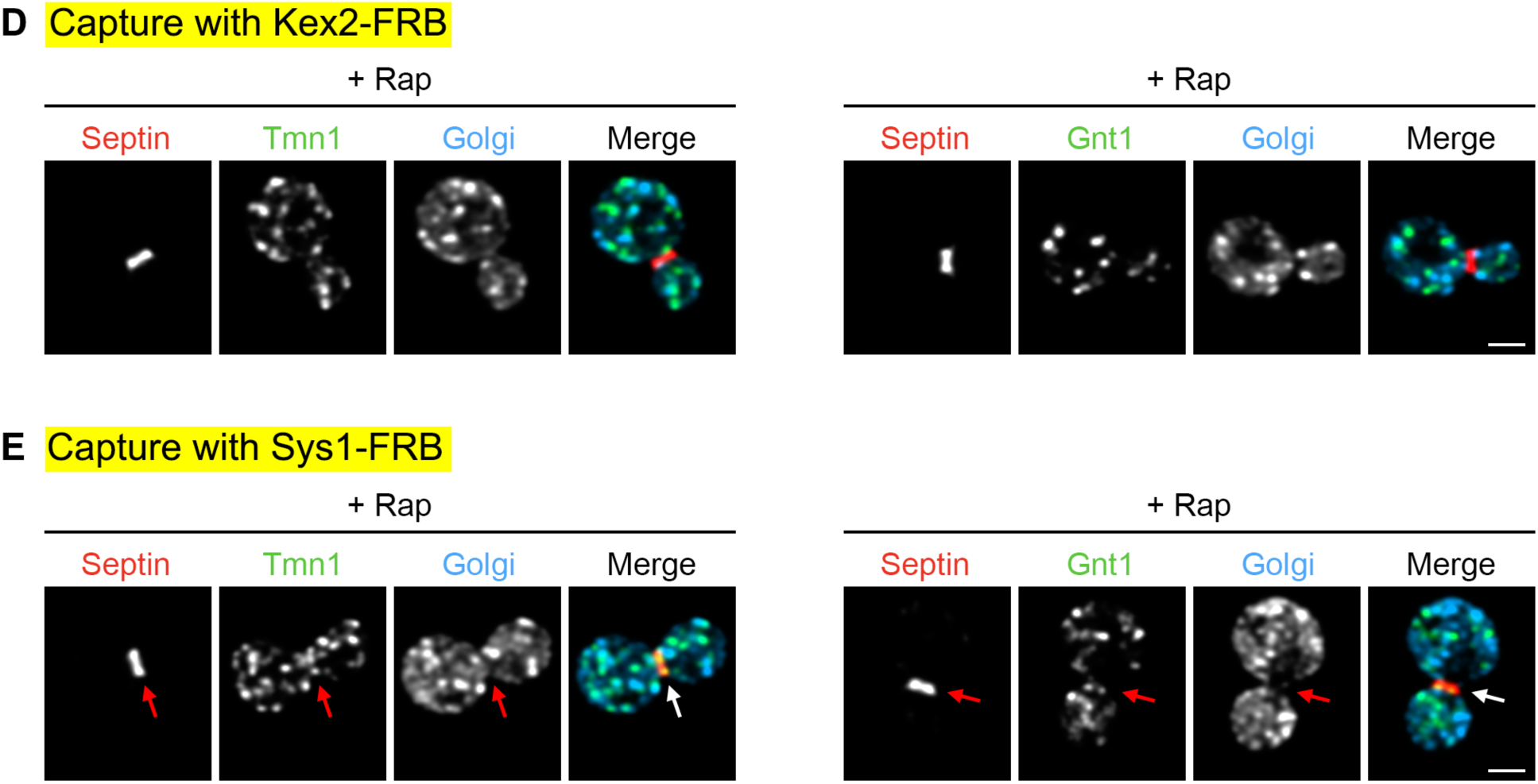
Supplementary data for Fig. 4. **(A)** Golgi maturation kinetics of GFP-tagged Vrg4 compared to HaloTag-labeled Sys1. Shown are normalized and averaged traces for 13 individual cisternae. **(B)** Kinetic traces for two representative cisternae illustrating variations in the relative arrival and departure times of GFP-Tmn1 versus Sys1-HaloTag. Plotted are normalized fluorescence intensities. **(C)** Kinetic traces for three representative cisternae illustrating variations in the relative arrival and departure times of Gnt1-HaloTag versus GFP-Tmn1. Plotted are normalized fluorescence intensities. **(D)** Representative images showing minimal capture with Kex2-FRB of GFP-tagged Tmn1 (green) or no capture with Kex2-FRB of GFP-tagged Gnt1 (green) by an FKBP-tagged septin (red) after treatment for 5 min with rapamycin. **(E)** Representative images showing capture with Sys1-FRB of GFP-tagged Tmn1 or Gnt1 (green) by an FKBP-tagged septin (red) after treatment for 5 min with rapamycin. Arrows indicate non-Golgi signal at the bud neck.

**Figure S4.**
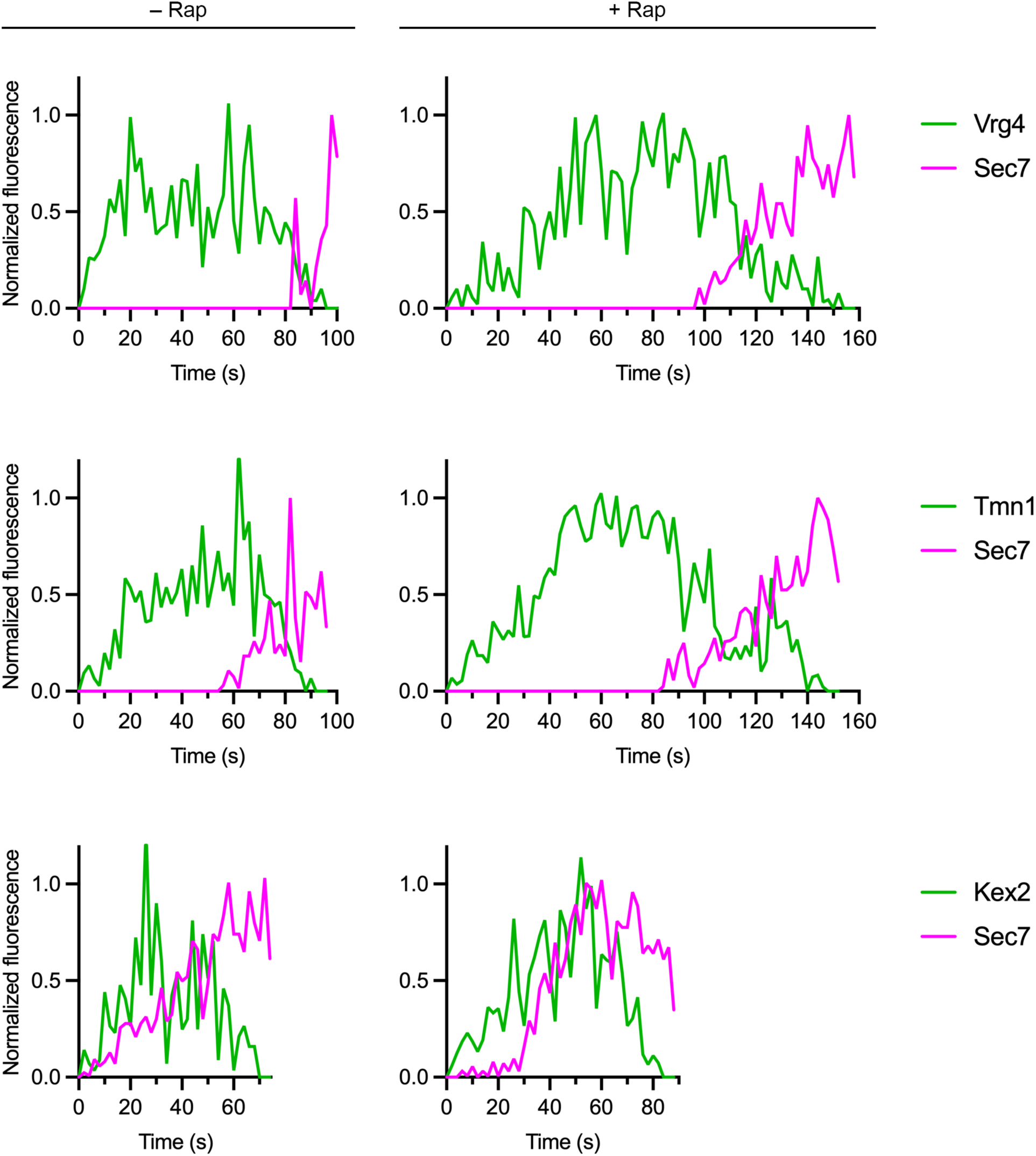
Supplementary data for Fig. 6. Kinetic traces for representative cisternae illustrate the residence times of GFP-tagged Vrg4, Tmn1, and Kex2 in Golgi cisternae either with normal COPI activity (“– Rap”) or with reduced COPI activity caused by brief rapamycin treatment (“+ Rap”). The analysis included partial companion traces of the late Golgi marker Sec7-mScarlet. Plotted are normalized fluorescence intensities.

**Figure S5.**
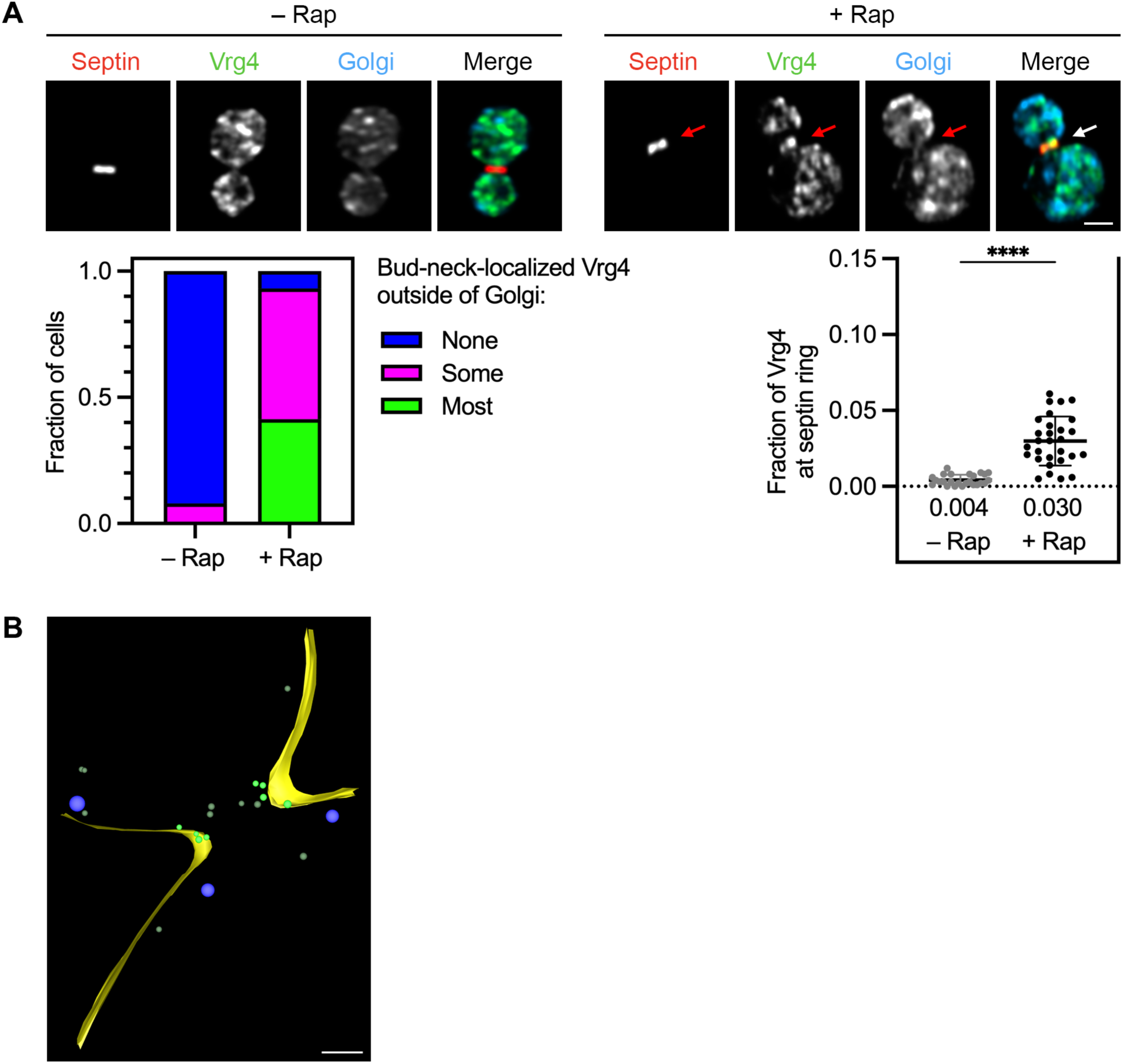
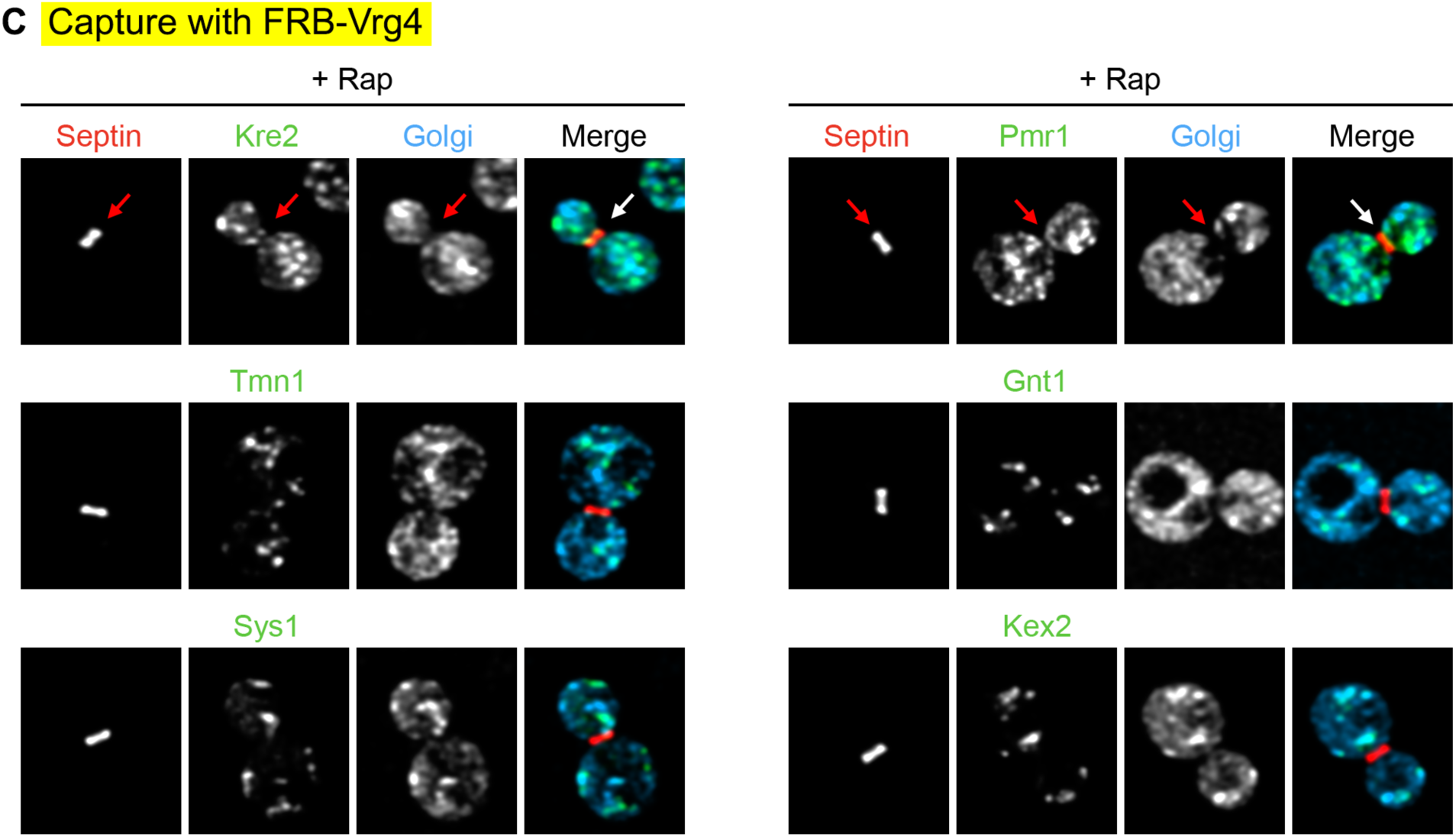
Supplementary data for Fig. 7. **(A)** Rapamycin-dependent capture by an FKBP-tagged septin (red) of FRB-GFP-Vrg4 (green). Arrows indicate non-Golgi signal at the bud neck after treatment for 5 min with rapamycin (“Rap”). At the lower left, capture of FRB-GFP-Vrg4 at the bud neck was quantified by assigning cells to categories as described in Fig. 1 C. At the lower right, capture of FRB-GFP-Vrg4 at the bud neck was quantified numerically as described in Fig. 1 C. ****, significant at P value <0.0001. **(B)** Cryo-ET of vesicles captured at the bud neck using FRB-Vrg4. A log-phase culture of cells expressing FRB-Vrg4 and Shs1-FKBP was treated with rapamycin for 5 min followed by cryopreservation and processing for cryo-ET. Shown is the model of a SIRT-reconstructed tomogram from a large budded cell. The full data set is shown in Video 6. A vesicle was counted as putatively captured if its membrane was no more than 66 nm from a point on the plasma membrane within 200 nm from the center of the bud neck. For the 6 non-rapamycin-treated cells examined, the number of vesicles meeting this criterion ranged from 0 to 2 (mean = 0.8). For the 6 rapamycin-treated cells examined, the number of putatively captured vesicles ranged from 2 to 7 (mean = 5.0). Scale bar, 250 nm. See Fig. 1 D for further details. **(C)** Representative images showing capture with FRB-Vrg4 of GFP-tagged Kre2 and Pmr1 (green) by an FKBP-tagged septin (red), minimal capture with FRB-Vrg4 of GFP-tagged Tmn1 and Sys1 (green) by an FKBP-tagged septin (red), and no capture with FRB-Vrg4 of GFP-tagged Gnt1 or Kex2 (green) by an FKBP-tagged septin (red), all after treatment for 5 min with rapamycin. Arrows indicate non-Golgi signal at the bud neck.

**Video 1.**
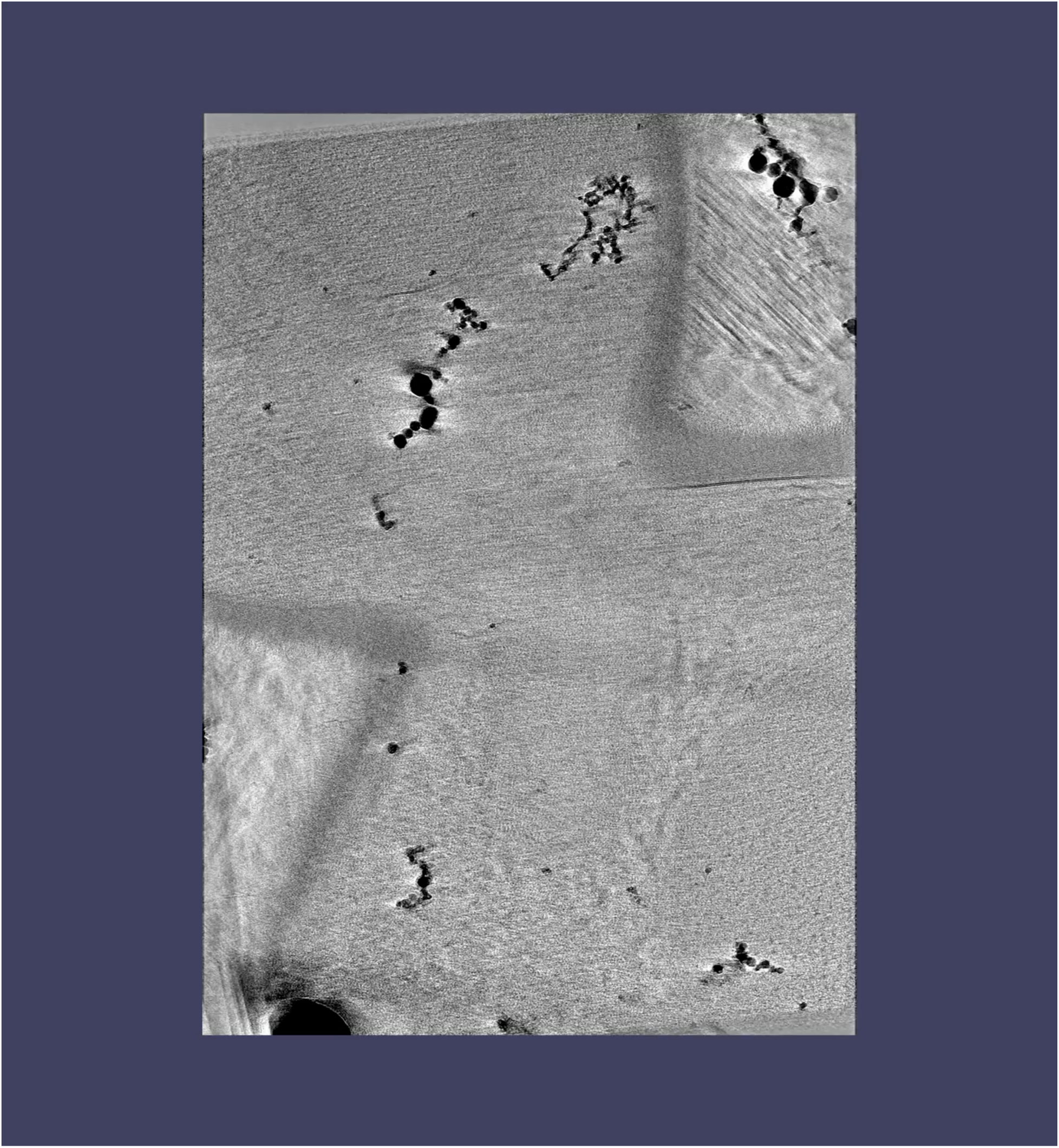
Tomographic sections and modeling of part of the bud neck region in a cell with Kex2-FRB-containing vesicles captured by an FKBP-tagged septin. A log-phase culture of cells expressing Kex2-FRB and Shs1-FKBP was treated for 5 min with rapamycin prior to cryopreservation and processing for cryo-ET. The first third of the video shows every fifth section of the SIRT-reconstructed tomogram. The second third of the video shows the same tomographic sections after contours were segmented to mark the cell cortex (yellow), a secretory vesicle (blue), putatively captured non-secretory vesicles (bright green), and other non-secretory vesicles (dull green). Also marked is a mitochondrion (cyan). The final third of the video shows a rotation of the tomographic model.

**Video 2.**
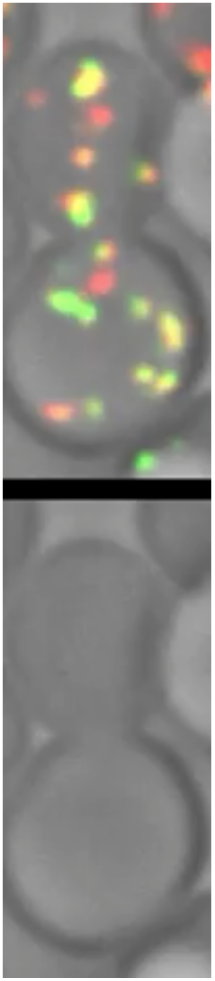
Representative 4D confocal movie of Sys1-HaloTag and Kex2-GFP. 3D z-stacks for the individual time points were average projected. The upper row shows the complete projections, and the lower row shows edited projections that include only the cisterna that was tracked. Intervals between frames are 2 s. The overlaid numbers in a subset of the frames represent the time in seconds after the cisterna that was tracked first became detectable. See Fig. S2 A for further details.

**Video 3.**
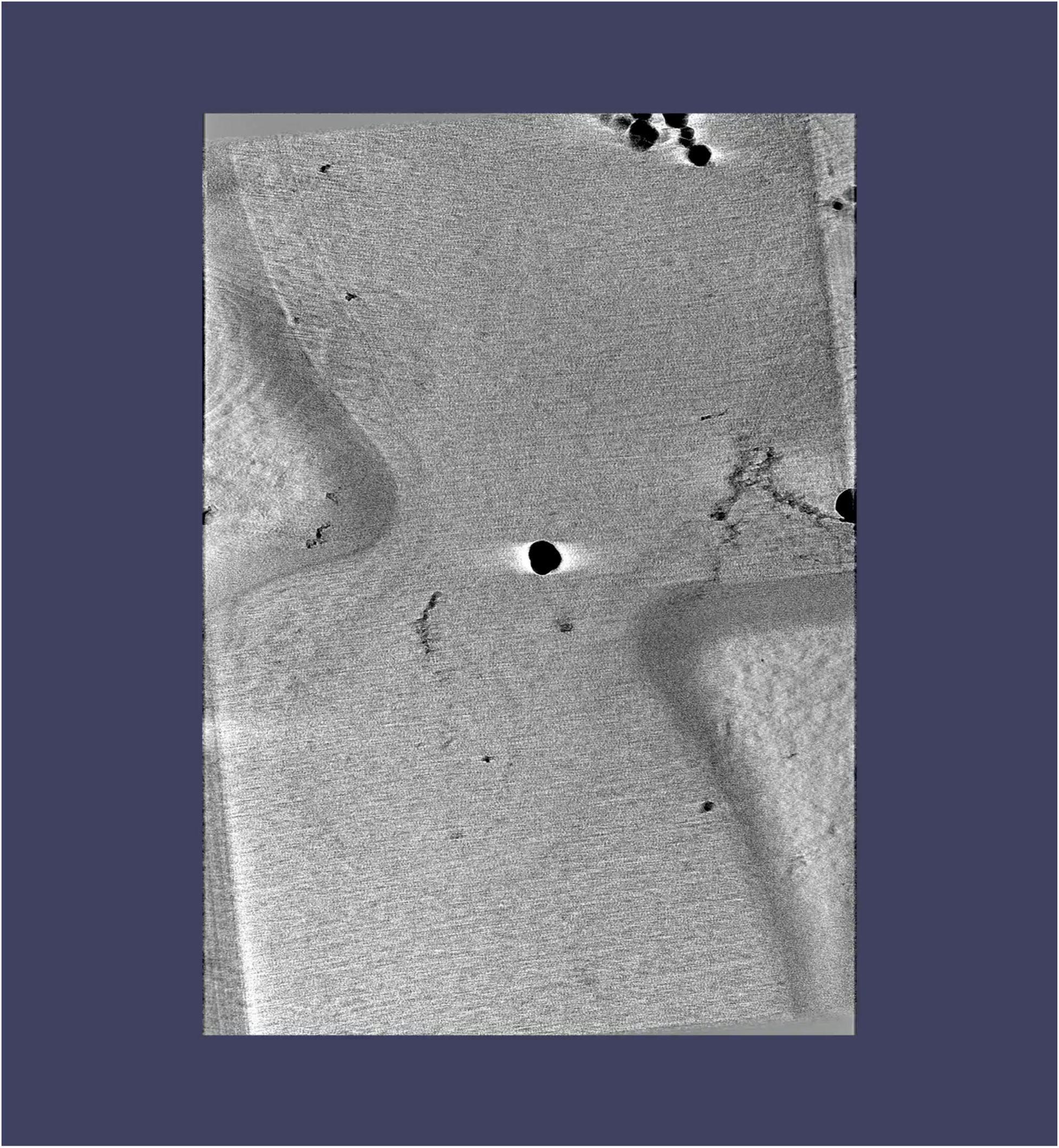
Tomographic sections and modeling of part of the bud neck region in a cell with Sys1-FRB-containing vesicles captured by an FKBP-tagged septin. A log-phase culture of cells expressing Sys1-FRB and Shs1-FKBP was treated for 5 min with rapamycin prior to cryopreservation and processing for cryo-ET. Further details are as in Video 1, except that the nuclear envelope is marked in magenta.

**Video 4.**
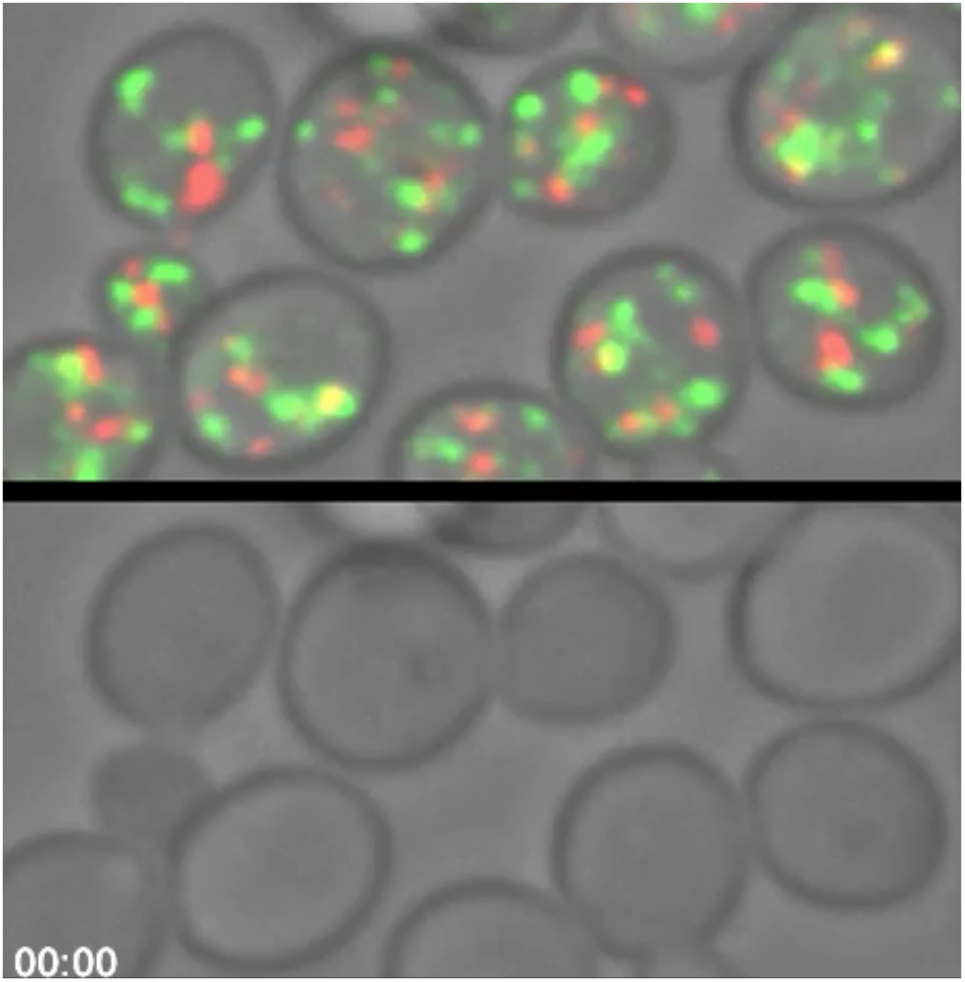
Representative 4D confocal movie of Vrg4 dynamics in Golgi cisternae with normal COPI activity. Intervals between frames are 2 s. This movie corresponds to the “– Rap” trace at the top of Fig. S4, with time zero in that trace corresponding to 2:22 for the numbering shown here. 3D z-stacks for the individual time points were average projected. The upper row shows the complete projections, and the lower row shows edited projections that include only the single cisterna that was tracked. Vrg4 is green and Sec7 is red.

**Video 5.**
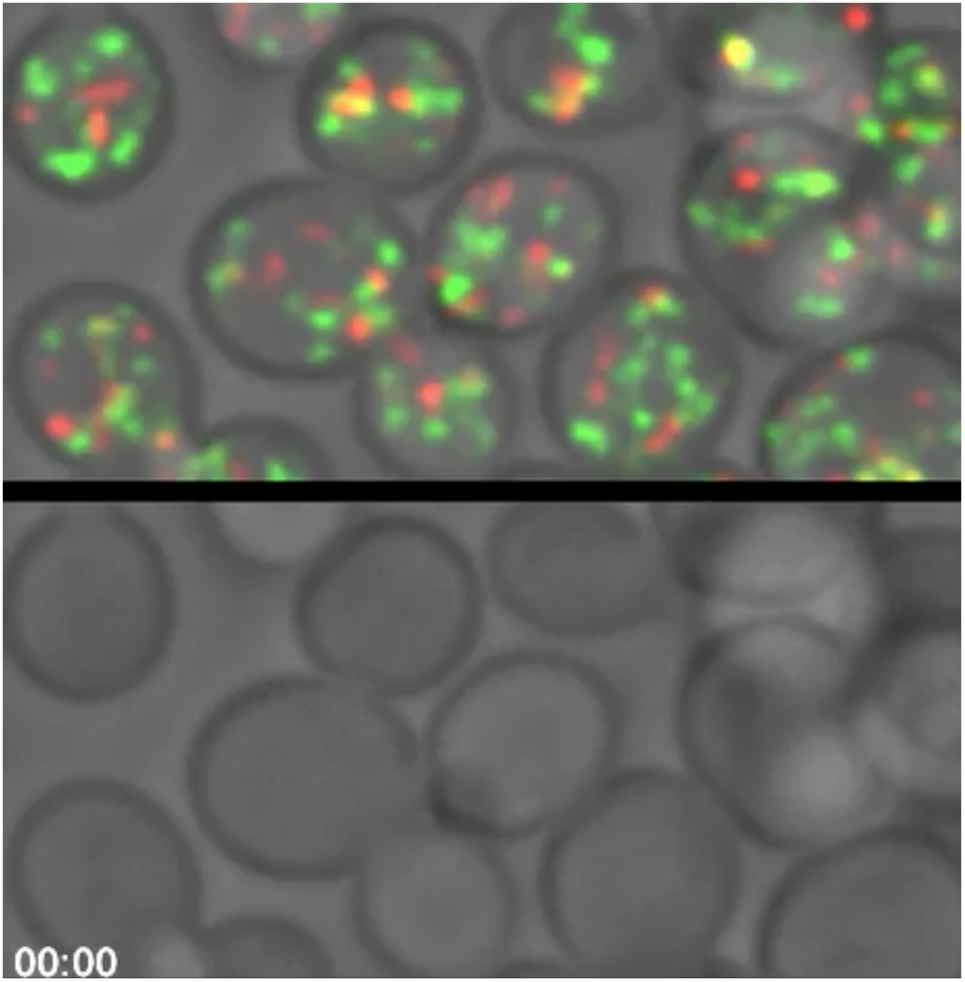
Representative 4D confocal movie of Vrg4 dynamics in Golgi cisternae with compromised COPI activity. Time zero in this movie is 4 min after rapamycin addition, and intervals between frames are 2 s. This movie corresponds to the “+ Rap” trace at the top of Fig. S4, with time zero in that trace corresponding to 2:56 for the numbering shown here. 3D z-stacks for the individual time points were average projected. The upper row shows the complete projections, and the lower row shows edited projections that include only the single cisterna that was tracked. Vrg4 is green and Sec7 is red.

**Video 6.**
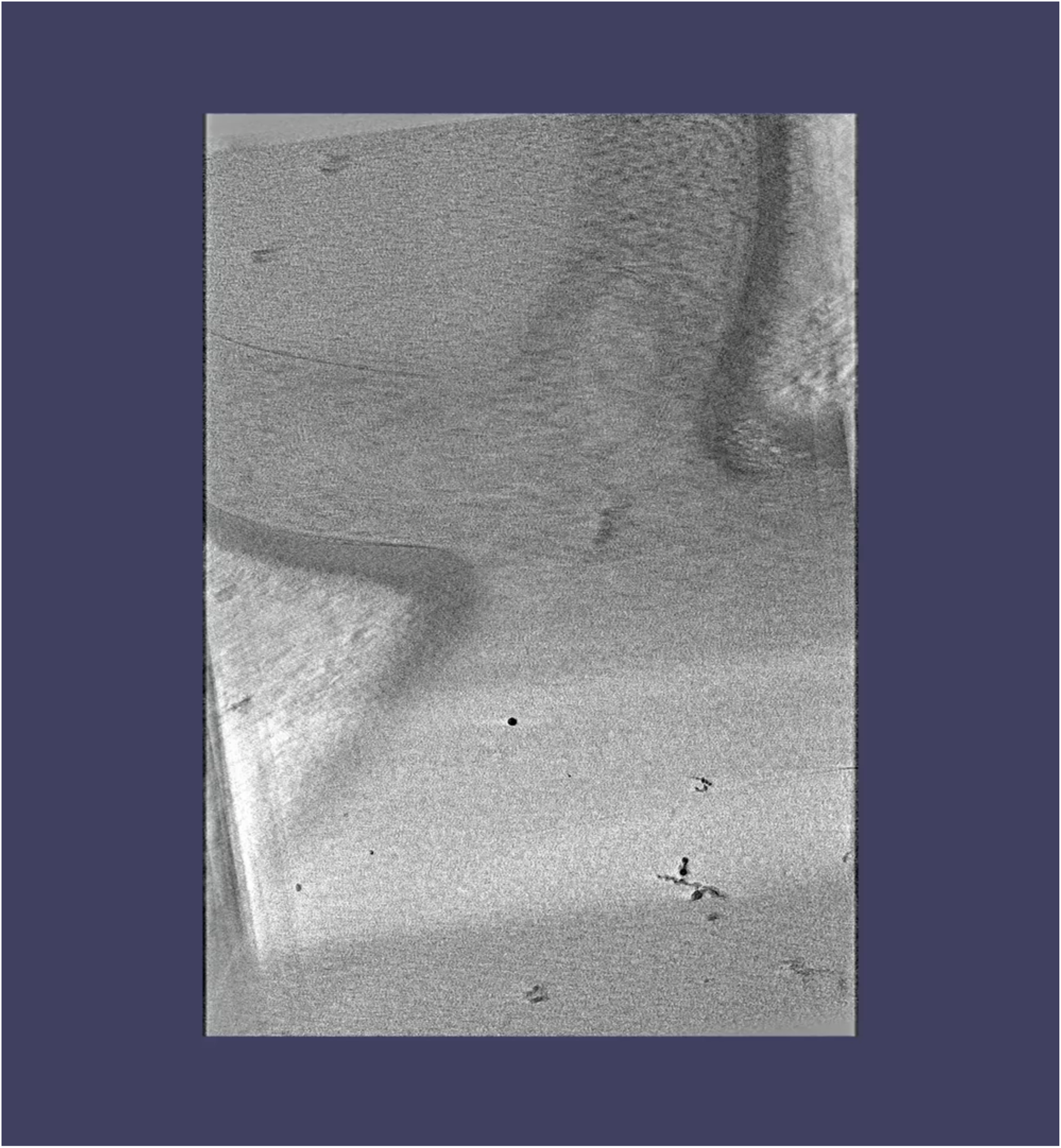
Tomographic sections and modeling of part of the bud neck region in a cell with FRB-Vrg4-containing vesicles captured by an FKBP-tagged septin. A log-phase culture of cells expressing FRB-Vrg4 and Shs1-FKBP was treated for 5 min with rapamycin prior to cryopreservation and processing for cryo-ET. Further details are as in Video 1.

## References

Abe, M., Y. Noda, H. Adachi, and K. Yoda. 2004. Localization of GDP-mannose transporter in the Golgi requires retrieval to the endoplasmic reticulum depending on its cytoplasmic tail and coatomer. J. Cell Sci. 117:5687–5696.

Abramson, J., J. Adler, J. Dunger, R. Evans, T. Green, A. Pritzel, O. Ronneberger, L. Willmore, A.J. Ballard, J. Bambrick, S.W. Bodenstein, D.A. Evans, C.C. Hung, M. O’Neill, D. Reiman, K. Tunyasuvunakool, Z. Wu, A. Žemgulytė, E. Arvaniti, C. Beattie, O. Bertolli, A. Bridgland, A. Cherepanov, M. Congreve, A.I. Cowen-Rivers, A. Cowie, M. Figurnov, F.B. Fuchs, H. Gladman, R. Jain, Y.A. Khan, C.M.R. Low, K. Perlin, A. Potapenko, P. Savy, S. Singh, A. Stecula, A. Thillaisundaram, C. Tong, S. Yakneen, E.D. Zhong, M. Zielinski, A. Žídek, V. Bapst, P. Kohli, M. Jaderberg, D. Hassabis, and J.M. Jumper. 2024. Accurate structure prediction of biomolecular interactions with AlphaFold 3. Nature. 630:493–500.

Antebi, A., and G.R. Fink. 1992. The yeast Ca^2+^-ATPase homologue, PMR1, is required for normal Golgi function and localizes in a novel Golgi-like distribution. Mol. Biol. Cell. 3:633–654.

Arab, M., T. Chen, and M. Lowe. 2024. Mechanisms governing vesicle traffic at the Golgi apparatus. Curr. Opin. Cell Biol. 88:102365.

Barlowe, C.K., and E.A. Miller. 2013. Secretory protein biogenesis and traffic in the early secretory pathway. Genetics. 193:383–410.

Barrero, J.J., E. Papanikou, J.C. Casler, K.J. Day, and B.S. Glick. 2016. An improved reversibly dimerizing mutant of the FK506-binding protein FKBP. Cell. Logist. 6:e1204848.

Bertin, A., M.A. McMurray, J. Pierson, L. Thai, K.L. McDonald, E.A. Zehr, G. García, 3rd, P. Peters, J. Thorner, and E. Nogales. 2012. Three-dimensional ultrastructure of the septin filament network in *Saccharomyces cerevisiae*. Mol. Biol. Cell. 23:423–432.

Bevis, B.J., A.T. Hammond, C.A. Reinke, and B.S. Glick. 2002. *De novo* formation of transitional ER sites and Golgi structures in *Pichia pastoris*. Nat. Cell Biol. 4:750–756.

Bindels, D.S., L. Haarbosch, L. van Weeren, M. Postma, K.E. Wiese, M. Mastop, S. Aumonier, G. Gotthard, A. Royant, M.A. Hink, and T.W.J. Gadella. 2017. mScarlet: a bright monomeric red fluorescent protein for cellular imaging. Nat. Methods. 14:53–56.

Casler, J.C., and B.S. Glick. 2020. A microscopy-based kinetic analysis of yeast vacuolar protein sorting. eLife. 9:e56844.

Casler, J.C., N. Johnson, A.H. Krahn, A. Pantazopoulou, K.J. Day, and B.S. Glick. 2022. Clathrin adaptors mediate two sequential pathways of intra-Golgi recycling. J. Cell. Biol. 221:e202103199.

Casler, J.C., E. Papanikou, J.J. Barrero, and B.S. Glick. 2019. Maturation-driven transport and AP-1-dependent recycling of a secretory cargo in the Golgi. J. Cell Biol. 218:1582–1601.

Cho, J.H., Y. Noda, and K. Yoda. 2000. Proteins in the early golgi compartment of *Saccharomyces cerevisiae* immunoisolated by Sed5p. FEBS Lett. 469:151–154.

Comet, M., P.M. Dijkman, R. Boer Iwema, T. Franke, S. Masiulis, R. Schampers, O. Raschdorf, F. Grollios, E.E.J. Pryor, and I. Drulyte. 2024. Tomo Live: an on-the-fly reconstruction pipeline to judge data quality for cryo-electron tomography workflows. Acta Crystallogr. D Struct. Biol. 80:247–258.

Daboussi, L., G. Costaguta, and G.S. Payne. 2012. Phosphoinositide-mediated clathrin adaptor progression at the *trans*-Golgi network. Nat. Cell Biol. 14:239–248.

Day, K.J., J.C. Casler, and B.S. Glick. 2018. Budding yeast has a minimal endomembrane system. Dev. Cell. 44:56–72.

Dürr, G., J. Strayle, R. Plemper, S. Elbs, S.K. Klee, P. Catty, D.H. Wolf, and H.K. Rudolph. 1998. The medial-Golgi ion pump Pmr1 supplies the yeast secretory pathway with Ca^2+^ and Mn^2+^ required for glycosylation, sorting, and endoplasmic reticulum-associated protein degradation. Mol. Biol. Cell. 9:1149–1162.

Farquhar, M.G., and H.-P. Hauri. 1997. Protein sorting and vesicular traffic in the Golgi apparatus. *In* The Golgi Apparatus. E.G. Berger and J. Roth, editors. Birkhäuser Verlag, Basel. 63–129.

Fegan, A., B. White, J.C. Carlson, and C.R. Wagner. 2010. Chemically controlled protein assembly: techniques and applications. Chem. Rev. 110:3315–3336.

Finger, F.P., T.E. Hughes, and P. Novick. 1998. Sec3p is a spatial landmark for polarized secretion in budding yeast. Cell. 92:559–571.

Fitzgerald, I., and B.S. Glick. 2014. Secretion of a foreign protein from budding yeasts is enhanced by cotranslational translocation and by suppression of vacuolar targeting. Microb. Cell Fact. 13:125.

Fuller, R.S., R.E. Sterne, and J. Thorner. 1988. Enzymes required for yeast prohormone processing. Annu. Rev. Physiol. 50:345–362.

Gan, L., C.T. Ng, C. Chen, and S. Cai. 2019. A collection of yeast cellular electron cryotomography data. Gigascience. 8:giz077.

Gaynor, E.C., T.R. Graham, and S.D. Emr. 1998. COPI in ER/Golgi and intra-Golgi transport: do yeast COPI mutants point the way? Biochim. Biophys. Acta. 1404:33–51.

Gilbert, P. 1972. Iterative methods for the three-dimensional reconstruction of an object from projections. J. Theor. Biol. 36:105–117.

Glick, B.S. 2025. Rethinking the yeast endomembrane system. *Subcell. Biochem*. Under editorial review.

Glick, B.S., T. Elston, and G. Oster. 1997. A cisternal maturation mechanism can explain the asymmetry of the Golgi stack. FEBS Lett. 414:177–181.

Glick, B.S., and V. Malhotra. 1998. The curious status of the Golgi apparatus. Cell. 95:883–889.

Glick, B.S., and A. Nakano. 2009. Membrane traffic within the Golgi stack. Annu. Rev. Cell Dev. Biol. 25:113–132.

Grimm, J.B., L. Xie, J.C. Casler, R. Patel, A.N. Tkachuk, N. Falco, H. Choi, J. Lippincott-Schwartz, T.A. Brown, B.S. Glick, Z. Liu, and L.D. Lavis. 2021. A general method to improve fluorophores using deterated auxochromes. JACS Au. 1:690–696.

Hagen, W.J.H., W. Wan, and J.A.G. Briggs. 2017. Implementation of a cryo-electron tomography tilt-scheme optimized for high resolution subtomogram averaging. J. Struct. Biol. 197:191–198.

Haruki, H., J. Nishikawa, and U.K. Laemmli. 2008. The anchor-away technique: rapid, conditional establishment of yeast mutant phenotypes. Mol. Cell. 31:925–932.

Huh, W.K., J.V. Falvo, L.C. Gerke, A.S. Carroll, R.W. Howson, J.S. Weissman, and E.K. O’Shea. 2003. Global analysis of protein localization in budding yeast. Nature. 425:686–691.

Iwase, M., J. Luo, E. Bi, and A. Toh-e. 2007. Shs1 plays separable roles in septin organization and cytokinesis in *Saccharomyces cerevisiae*. Genetics. 177:215–229.

Johnson, N., and B.S. Glick. 2019. 4D microscopy of yeast. J. Vis. Exp.:10.3791/58618.

Kim, J.J., Z. Lipatova, and N. Segev. 2016. Regulation of Golgi cisternal progression by Ypt/Rab GTPases. Dev. Cell. 36:440–452.

Kremer, J.R., D.N. Mastronarde, and J.R. McIntosh. 1996. Computer visualization of three-dimensional image data using IMOD. J. Struct. Biol. 116:71–76.

Kunz, J., U. Schneider, M. Deuter-Reinhard, N.R. Movva, and M.N. Hall. 1993. Target of rapamycin in yeast, TOR2, is an essential phosphatidylinositol kinase homolog required for G1 progression. Cell. 73:585–596.

Losev, E., C.A. Reinke, J. Jellen, D.E. Strongin, B.J. Bevis, and B.S. Glick. 2006. Golgi maturation visualized in living yeast. Nature. 441:1002–1006.

Lussier, M., A.M. Sdicu, T. Ketela, and H. Bussey. 1995. Localization and targeting of the *Saccharomyces cerevisiae* Kre2p/Mnt1p (alpha)1,2-mannosyltransferase to a *medial*-Golgi compartment. J. Cell Biol. 131:913–927.

Mari, M., W.J. Geerts, and F. Reggiori. 2014. Immuno- and correlative light microscopy-electron tomography methods for 3D protein localization in yeast. Traffic. 15:1164–1178.

Matsuura-Tokita, K., M. Takeuchi, A. Ichihara, K. Mikuriya, and A. Nakano. 2006. Live imaging of yeast Golgi cisternal maturation. Nature. 441:1007–1010.

McDonold, C.M., and J.C. Fromme. 2014. Four GTPases differentially regulate the Sec7 Arf-GEF to direct traffic at the *trans*-Golgi network. Dev. Cell. 30:759–767.

Mowbrey, K., and J.B. Dacks. 2009. Evolution and diversity of the Golgi body. FEBS Lett. 583:3738–3745.

Myers, M.D., and G.S. Payne. 2013. Clathrin, adaptors and disease: insights from the yeast *Saccharomyces cerevisiae*. Front. Biosci. 18:862–891.

Ong, K., C. Wloka, S. Okada, T. Svitkina, and E. Bi. 2014. Architecture and dynamic remodelling of the septin cytoskeleton during the cell cycle. Nat. Commun. 5:5698.

Pantazopoulou, A., and B.S. Glick. 2019. A kinetic view of membrane traffic pathways can transcend the classical view of Golgi compartments. Front. Cell Dev. Biol. 7:153.

Papanikou, E., K.J. Day, J. Austin, 2nd, and B.S. Glick. 2015. COPI selectively drives maturation of the early Golgi. eLife. 4:e13232.

Pruyne, D., A. Legesse-Miller, L. Gao, Y. Dong, and A. Bretscher. 2004. Mechanisms of polarized growth and organelle segregation in yeast. Annu. Rev. Cell Dev. Biol. 20:559–591.

Rabouille, C., and J. Klumperman. 2005. Opinion: The maturing role of COPI vesicles in intra-Golgi transport. Nat. Rev. Mol. Cell Biol. 6:812–817.

Robinson, M.S., R. Antrobus, A. Sanger, A.K. Davies, and D.C. Gershlick. 2024. The role of the AP-1 adaptor complex in outgoing and incoming membrane traffic. J. Cell Biol. 223:e202310071.

Rossanese, O.W., J. Soderholm, B.J. Bevis, I.B. Sears, J. O’Connor, E.K. Williamson, and B.S. Glick. 1999. Golgi structure correlates with transitional endoplasmic reticulum organization in *Pichia pastoris* and *Saccharomyces cerevisiae*. J. Cell Biol. 145:69–81.

Rothstein, R. 1991. Targeting, disruption, replacement, and allele rescue: integrative DNA transformation in yeast. Methods Enzymol. 194:281–301.

Sahu, P., A. Balakrishnan, R. Di Martino, A. Luini, and D. Russo. 2022. Role of the *mosaic* cisternal maturation machinery in glycan synthesis and oncogenesis. Front. Cell Dev. Biol. 10:842448.

Schimmöller, F., E. Díaz, B. Mühlbauer, and S.R. Pfeffer. 1998. Characterization of a 76 kDa endosomal, multispanning membrane protein that is highly conserved throughout evolution. Gene. 216:311–318.

Schmitz, K.R., J. Liu, S. Li, T.G. Setty, C.S. Wood, C.G. Burd, and K.M. Ferguson. 2008. Golgi localization of glycosyltransferases requires a Vps74p oligomer. Dev. Cell. 14:523–534.

Schüller, C., Y.M. Mamnun, H. Wolfger, N. Rockwell, J. Thorner, and K. Kuchler. 2007. Membrane-active compounds activate the transcription factors Pdr1 and Pdr3 connecting pleiotropic drug resistance and membrane lipid homeostasis in *Saccharomyces cerevisiae*. Mol. Biol. Cell. 18:4932–4944.

Singer-Krüger, B., R. Frank, F. Crausaz, and H. Riezman. 1993. Partial purification and characterization of early and late endosomes from yeast. Identification of four novel proteins. J. Biol. Chem. 268:14376–14386.

Siniossoglou, S., S.Y. Peak-Chew, and H.R. Pelham. 2000. Ric1p and Rgp1p form a complex that catalyses nucleotide exchange on Ypt6p. EMBO J. 19:4885–4894.

Tojima, T., Y. Suda, M. Ishii, K. Kurokawa, and A. Nakano. 2019. Spatiotemporal dissection of the *trans*-Golgi network in budding yeast. J. Cell Sci. 132: jcs231159.

Tojima, T., Y. Suda, N. Jin, K. Kurokawa, and A. Nakano. 2024. Spatiotemporal dissection of the Golgi apparatus and the ER-Golgi intermediate compartment in budding yeast. eLife. 13:e92900.

Tu, L., W.C. Tai, L. Chen, and D.K. Banfield. 2008. Signal-mediated dynamic retention of glycosyltransferases in the Golgi. Science. 321:404–407.

Valbuena, F.M., I. Fitzgerald, R.L. Strack, N. Andruska, L. Smith, and B.S. Glick. 2020. A photostable monomeric superfolder green fluorescent protein. Traffic. 21:534–544.

Welch, L.G., S.Y. Peak-Chew, F. Begum, T.J. Stevens, and S. Munro. 2021. GOLPH3 and GOLPH3L are broad-spectrum COPI adaptors for sorting into intra-Golgi transport vesicles. J. Cell Biol. 220:e202106115.

Wong, E.D., S.R. Miyasato, S. Aleksander, K. Karra, R.S. Nash, M.S. Skrzypek, S. Weng, S.R. Engel, and J.M. Cherry. 2023. Saccharomyces genome database update: server architecture, pan-genome nomenclature, and external resources. Genetics. 224:iyac191.

Wong, M., and S. Munro. 2014. Membrane trafficking. The specificity of vesicle traffic to the Golgi is encoded in the golgin coiled-coil proteins. Science. 346:1256898.

Woo, C.H., C. Gao, P. Yu, L. Tu, Z. Meng, D.K. Banfield, X. Yao, and L. Jiang. 2015. Conserved function of the lysine-based KXD/E motif in Golgi retention for endomembrane proteins among different organisms. Mol. Biol. Cell. 26:4280–4293.

Wooding, S., and H.R.B. Pelham. 1998. The dynamics of Golgi protein traffic visualized in living yeast cells. Mol. Biol. Cell. 9:2667–2680.

Yoko-o, T., C.A.R. Wiggins, J. Stolz, S.Y. Peak Chew, and S. Munro. 2003. An *N*-acetylglucosaminyltransferase of the Golgi apparatus of the yeast *Saccharomyces cerevisiae* that can modify N-linked glycans. Glycobiology. 13:581–589.

